# The plant circadian clock gene *LHY* influences *Medicago truncatula* nodulation

**DOI:** 10.1101/2021.03.22.435813

**Authors:** M Achom, P Roy, B Lagunas, R Bonyadi-Pour, AJ Pardal, L Baxter, B Richmond, N Aschauer, E Fletcher, E Picot, M Rowson, J Blackwell, C Rich-Griffin, KS Mysore, J Wen, S Ott, IA Carré, ML Gifford

## Abstract

Legumes house nitrogen-fixing endosymbiotic rhizobia in specialized polyploid cells within root nodules, which are factories of metabolic activity. We discovered that the circadian clock-associated transcriptional factor LATE ELONGATED HYPOCOTYL (LHY) affects nodulation in *Medicago truncatula*. By carrying out expression analysis of transcripts over time in nodules we found that the clock enables coordinated control of metabolic and regulatory processes linked to nitrogen fixation. Rhythmic transcripts in root nodules include a subset of Nodule-specific Cysteine Rich peptides (NCRs) that have the LHY-bound conserved Evening Element in their promoters. Until now, studies have suggested that NCRs act to regulate bacteroid differentiation and keep the rhizobial population in check. However, these conclusions came from the study of a few members of this very large gene family that has complex diversified spatio-temporal expression. We suggest that rhythmic expression of NCRs may be important for temporal coordination of bacterial activity with the rhythms of the plant host, in order to ensure optimal symbiosis.

**Highlights:**

- The circadian clock-associated transcriptional factor LATE ELONGATED HYPOCOTYL (LHY) impacts on successful *Medicago truncatula*-rhizobial symbiosis
- The plant clock coordinates rhythmic patterns of metabolic and regulatory activity in nodules and drives rhythmic expression of a subset of Nodule-specific Cysteine Rich (NCR) genes.
- Rhythmic expression of NCRs may be important for temporal coordination of bacterial activity with plant host rhythms to ensure optimal symbiosis.

## Introduction

In plants, animals and microbes, many aspects of physiology, metabolism and development exhibit 24-hour rhythmicity controlled by a circadian clock. Under natural day-night conditions, the circadian clock is synchronized to light-dark and temperature cycles and enables anticipation of predictable daily changes in the environment. Rhythmicity is particularly pervasive in plants. In the model plant *Arabidopsis thaliana*, about 30% of genes are expressed rhythmically in constant light, and up to 90% under at least some cycling environmental conditions (1). Many aspects of metabolism are rhythmic, including photosynthetic carbon assimilation, nitrogen and sulphur metabolism (2), and appropriate timing of starch utilization is known to ensure optimal growth (3). The circadian clock also impacts plant productivity and health by modulating interactions with microorganisms. Plants show different levels of resistance to fungal and bacterial pathogens depending on the time of infection (4-7), and bacterial infections have been found to alter plant circadian regulation in order to attenuate immune responses (8). Plant circadian rhythms also influence the composition of rhizosphere microbial communities, and impeded circadian clock function in the plant host results in the recruitment of a different root microbiome, with consequences for plant health (9, 10).

The mechanism of the plant circadian clock has been studied extensively and shown to consist of a small gene network comprising multiple transcriptional feedback loops (11). In *A. thaliana*, a pair of closely related MYB transcription factors, Late Elongated Hypocotyl (LHY) and Circadian Clock Associated (CCA1), are expressed in the morning and act to repress the expression of other clock components, by binding to a DNA sequence motif in their promoters known as the Evening Element or EE (AAATATCT/AGATATTT) (2). As *LHY/CCA1* expression declines, a set of pseudo-response regulators (*PRR9, PRR7, PRR5* and *PRR1*, also known as *TOC1*) are expressed as sequential waves during the day and early evening, and act to repress expression of *LHY* and *CCA1* till the following dawn. A third set of proteins, composed of LUX ARRYTHMO (LUX), EARLY FLOWERING (ELF3) and ELF4, is expressed at dusk and forms an ‘Evening Complex’. There is evidence that a similar mechanism operates in roots, although whereas the leaf clock is primarily synchronized to diurnal light-dark cycles, the root clock is thought to be entrained by shoot-derived signals (12-14). Clock components are conserved in both monocot and dicot crops, and have been linked to important agronomic traits including growth and flowering time (15). Homologues of *A. thaliana* clock genes have been identified in most legumes including soybean (*Glycine max*) cow pea (*Vigna unguiculata*) and garden pea (*Pisum sativum*) (16-19). While LHY and CCA1 have largely redundant function in the central oscillator of *A. thaliana*, a single orthologue of these proteins is present in *Medicago truncatula*, termed *MtLHY* (20).

Altered function of the soybean circadian clock through overexpression of a light-signalling component has been seen to lead to grain yield increases (21). However, there is a lack of information about the impact of the circadian clock on legume symbioses with nitrogen-fixing rhizobia. This is important, because this symbiosis contributes to the nitrogen nutrition of the plant which increases plant growth, reducing the need for synthetic nitrogen fertilisers while also improving soil health. During nodulation, rhizobia are accommodated in specialized lateral root organs called nodules. Formation of nodules is initiated following recognition, by host plant LysM receptors, of Nod factors (NF) released from rhizobial bacteria. This leads to activation of calcium oscillations, then transcriptional responses that enable controlled cell division for nodule formation, and rhizobial entry via an infection thread. Within nodules, rhizobia inhabit an intracellular compartment derived from host cell membranes, called the symbiosome. They proliferate and differentiate into nitrogen-fixing bacteroids, which convert atmospheric di-nitrogen into a plant-accessible form such as ammonium that the host plant will incorporate into its own nitrogen metabolism. In exchange for the fixed nitrogen, the bacteria benefits from host-supplied carbon and other nutrients (22, 23). The evolution of nodulation in legumes has been greatly shaped by a whole genome duplication event approximately 58 million years ago (MYA), resulting in amplified, rearranged gene families and retention of paralogous genes (24). Prominent amongst these is the Nodule Cysteine-Rich (NCR) gene family of small secreted peptides that are highly specific to nodules (25). Except for some Aeschynomene species from the relatively ancient dalbergoid lineage, NCRs are exclusively found in the Inverted Repeat-Lacking Clade (IRLC) of legumes which includes the model plant *M. truncatula* and many agriculturally important crops such as alfalfa, clovers, lentils, chickpea, garden pea and fava beans (26). Only a few NCRs have been characterized in detail so far, but a picture is emerging of the importance of functional diversity for this gene family (25). The diverse spatio-temporal expression profiles of NCRs (27-29), high level of expression specificity across nodules (30, 31), and variation in amino acid sequence and isoelectric points (32), could enable this functional variation. Recent findings show that a subset of NCRs is regulated by nitrogen availability and by autoregulation of nodulation, suggesting an additional role for NCRs in controlling nodule development depending on cues from the environment (33).

Here we show that in *M. truncatula* nodules, disrupted circadian rhythmicity through loss of function of the core clock gene *LHY* results in reduced nodulation, suggesting that the circadian clock may impact on plant-rhizobia interactions in nodules. We investigate potential mechanisms through analysis of the rhythmic transcriptome in nodules and reveal circadian control of a subset of NCR genes through EE motifs in their promoters. We suggest that circadian regulation of NCR gene expression in nodules may play a role to ensure temporal coordination of bacterial activity with the rhythms of the plant host. Optimising the timing of nodule-specific nitrogen fixation-regulatory peptides may allow improvement of nitrogen fixation without altering any aboveground clock features. This may represent an interesting target for sustainable agriculture of legume crops.

## Results

### *Loss of LHY* function disrupts circadian rhythms and impairs nodulation in *M. truncatula*

In *A. thaliana*, LHY and CCA1 function as transcriptional repressors (34, 35) and have a largely redundant function in the central oscillator of *A. thaliana*. In *M. truncatula* a single orthologue of these proteins is present (Medtr7g118330) that shares 36.6% identity at the amino acid level to *AtCCA1* (At2g46830) and 44.2% identity at the amino acid level to *AtLHY* (At1g01060) (Fig. 1A), thus we call it *MtLHY*. In order to test the importance of *MtLHY* in the regulation of nodulation, we isolated and characterized two mutants with reduced *MtLHY* expression from the Noble collection of retrotransposon insertion (*Tnt1*) lines (Fig. 1B). The *lhy-1* mutant contained an insertion upstream of the translational start site, likely in the promoter of *LHY*, and exhibited strongly reduced expression of the *LHY* transcript (20% of WT levels; Fig. 1C; Table S1). The second mutant, *lhy-2*, had an insertion at the end of the second-last exon and negligible expression of the *LHY* transcript (Fig. 1C; Table S1).

**Fig. 1.**
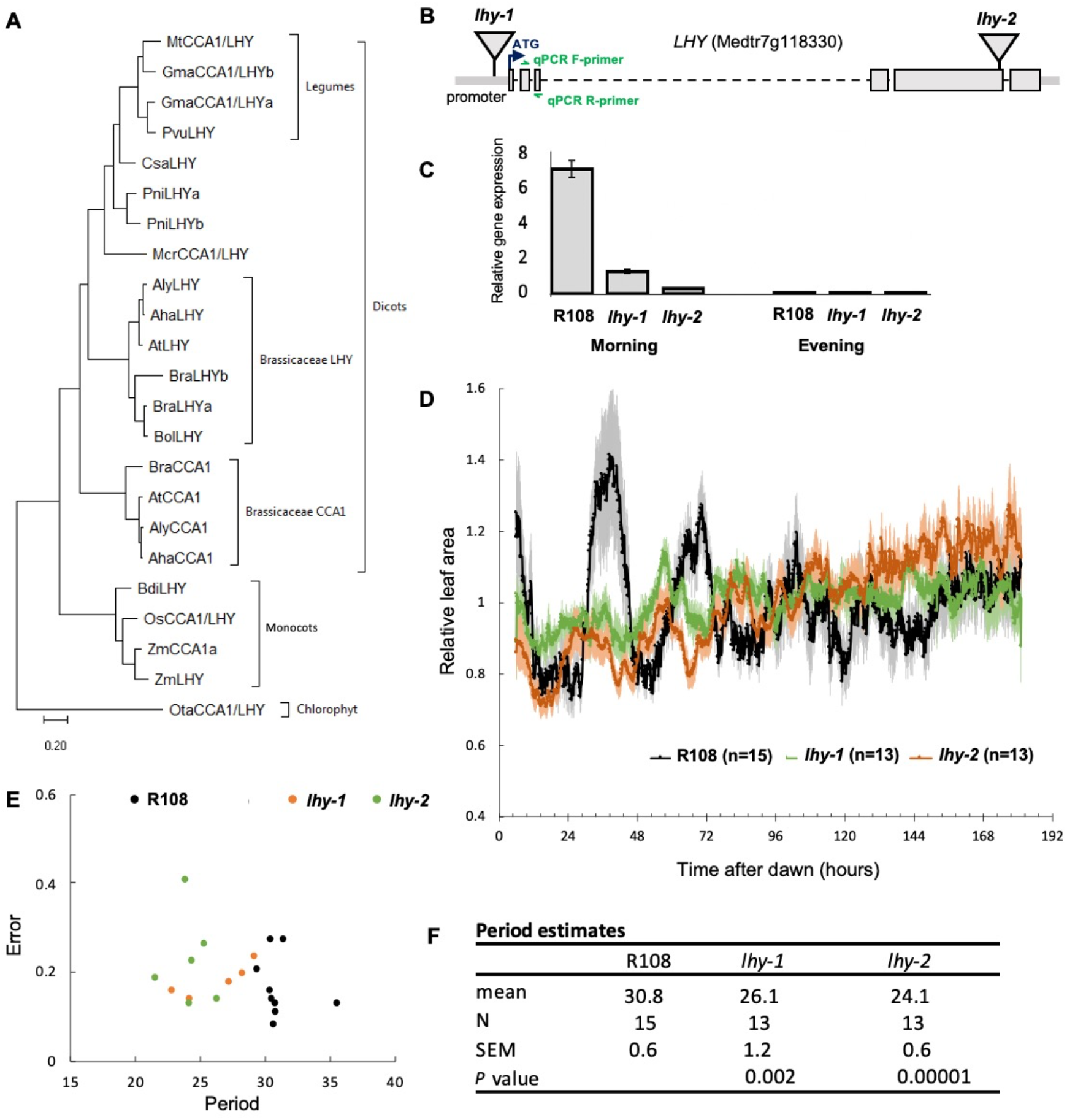
Loss of *M. truncatula LHY* expression affects plant rhythmicity and nodulation. (A) Phylogenetic analysis of CCA1/LHY homologues in Plantae and Chlorophyta; Aha, *Arabidopsis helleri*; Aly, *Arabidopsis lyrata*; At, *Arabidopis thaliana;* Bdi, *Brachypodium distachyon*; Bol, *Brassica oleracea*; Bra, *Brassica rapa*; Csa, *Castanea sativa*; Gma, *Glycine max*; Mcr, *Mesembryanthemum crystallinum*, Mt, *Medicago truncatula*; Os, *Oryza sativa*; Ot, *Ostreococcus tauri*; Pni, *Populus nigra*; Pvu, *Phaseolus vulgaris*; Zm, *Zea mays;* see Table S1 for gene IDs. (B) Location of *lhy-1* and *lhy-2* insertions in the *LHY* gene. (C) Relative expression of *MtLHY* in WT and mutant plant leaves in the morning and evening periods. (D) Disrupted leaf movement rhythms in *lhy* mutants in constant light; the experiment was repeated twice, then data from both biological replicates was combined. (E, F) Period analysis between 24 and 96 hours in constant light, for data shown in panel D. Period estimates shown correspond to leaf movement traces where the period analysis returned high quality fits to the data. Each data point in panel E corresponds to a period estimate for an individual leaf. Error values range between 0 and 1 and high error values are indicative of poor fit or weak rhythmicity. *P*-values in panel F are for t-test comparisons between wild-type and mutant means.

We examined the rhythmicity of our *lhy* mutants by measuring the rhythmic opening and closing of the first true leaf. Plants were grown for 10 days under 12L12D cycles then imaged over 7 days in constant light. The wild-type R108 showed sustained rhythmicity for over a week in constant light, with a period length of approximately 30 hours. In contrast, both mutant alleles exhibited leaf movement rhythms with shorter free-running periods (approximately 26 hours) and a much lower amplitude, then became arrhythmic after 120 hours (Fig. 1D-F). Leaf opening in the mutants occurred 6 hours early in the first day following transfer to constant light, indicating that both mutations resulted in a large phase advance in constant light (Fig. 1E-F). These results demonstrated that loss of *LHY* function alters the function of the clock in *M. truncatula*.

In order to assess the effect of *lhy* mutations on nodulation, we inoculated three biological replicates of *lhy-1, lhy-2* and R108 seedlings with the high-efficiency rhizobial symbiont *Sinorhizobium meliloti* WSM1022. When grown under 16L8D, both mutants had lower nodule weight and lower dry shoot weight than the wild-type (Fig. 2A-C; Table S1). Interestingly, our data shows that *lhy* yield is similar to WT when not inoculated with rhizobia, but reduced when plants are inoculated with rhizobia, suggesting that the reduced yield in the mutant is largely due to disrupted nodulation (Fig. 2A-C). These observations indicate that normal function of *LHY* is required for optimal nodulation and plant yield.

**Figure 2.**
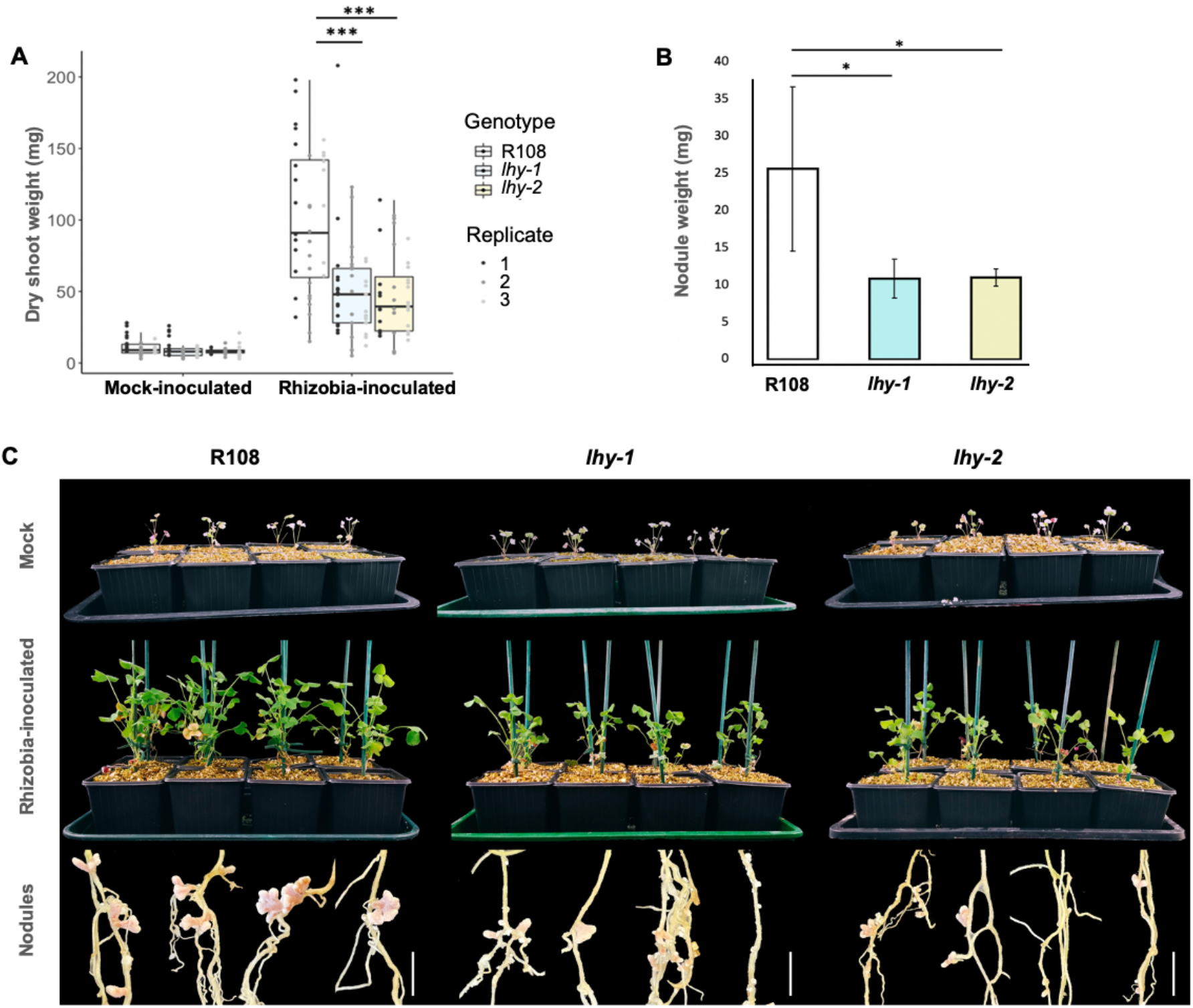
Loss of *M. truncatula LHY* expression affects nodulation under 16L8d cycles. (A) Plants have a similar dry aboveground weight phenotype in the absence of rhizobial inoculation, but with inoculation the *lhy* mutants have reduced dry weight; boxplots with individual replicate data; n=24; \*\*\**P*<0.005. (B) Reduced nodule weight for *lhy-1* and *lhy-2* compared to WT R108; n=24; \**P*<0.05. (C) Images of six week old mock (top row) or *Sinorhizobium meliloti* WSM1022-inoculated plants grown in perlite-vermiculate pots showing reduced growth (middle row) and nodule size (lower row) for the *lhy* mutants. See Table S1 for all values and analyses.

### Gene expression within nodules shows the presence of a functional nodule clock

We then asked what processes involved in nodule function might be under circadian regulation by carrying out a timecourse analysis of the rhythmic transcriptome. Plants were grown under diurnal light-dark cycles (12L12D) for 40 days in order to entrain their circadian clocks, then transferred to constant light to test for persistence of rhythms in the absence of environmental time cues. Nodules were sampled every 3 hrs for the first 24-hours in constant light, then every 6 hrs for another 24 hours (Fig. 3A), and changes in gene expression were analyzed by RNA-seq (Table S2). After normalization and mapping of reads to the *M. truncatula* genome (4.0v1) we were able to determine expression levels for 61,510 transcripts out of the 62,319 protein-coding transcripts in the genome (98.7%). We then used Metacycle analysis (36) to identify transcripts that oscillate with a period of approximately 24-hours. This identified 2,832 transcripts with rhythmic behaviour in constant light (*P*<0.05) (Fig. 3B), i.e about 5 % of the transcriptome. Hierarchical clustering identified four broad groups of genes (Fig. 3C-F). The majority of rhythmic genes peaked around dawn and were found in clusters 1 and 4, peaking 24 and 21 hours after dawn, respectively. These morning and late-night clusters contained 811 and 1,458 transcripts, respectively, whereas the morning and evening clusters (clusters 2 and 3) contained only 242 and 321 transcripts. This contrasted with previous observations in *A. thaliana* whole plants, that the majority of cycling genes peak either before dawn or dusk (1).

**Fig. 3.**
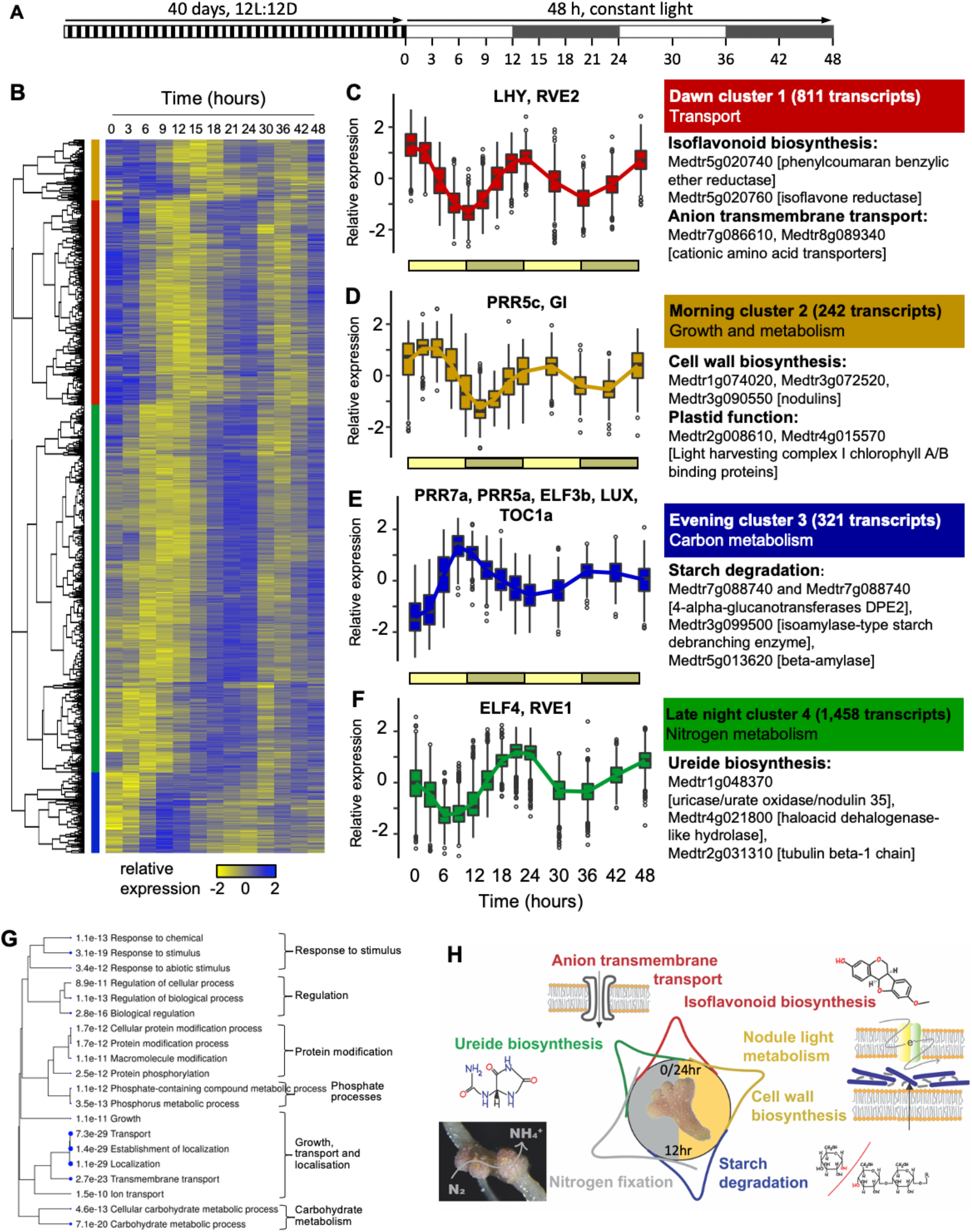
Oscillating expression of the nodule clock. (A) Experimental design for 48 hour timecourse, with sampling points (B) Heatmap of expression values of the 2,832 transcripts with a rhythmic pattern (*p*<0.05); see Table S2. (C-F) There are four clusters whose expression peaks at: dawn (red, C), morning (yellow, D), evening (blue, E), late night (green, F); overrepresented GO terms, pathways or protein domains and the clock-related genes (see Table S3) within each cluster are shown. (G) Hierarchical clustering representing correlation among significantly enriched (top 20) biological processes in rhythmic transcripts. Enrichment *P*-values (Bonferroni-corrected for multiple testing) are shown for each biological process and represented by blue dots, where larger dots are more highly significant; see Table S4. (H) Summary of changes over a 24h period (12L12D) in nodules.

We first examined the expression patterns of clock-associated genes in nodules; Fig. 3C-F; Table S3. Putative orthologs of all *A. thaliana* circadian clock genes are present within the *M. truncatula* genome (Table S3). *LHY* was identified in cluster 1 and peaked at dawn. Q-PCR analyses confirmed this observation, and showed that temporal patterns of *LHY* expression were similar in roots and nodules, but preceded expression in shoots by several hours in constant light (Fig. S1). *PRR5* homologues were expressed in consecutive waves, with *PRR5c* and *PRR5a* peaking 3 and 9 hours after dawn, and *PRR7a* and *TOC1* peaking at dusk (Fig. S2). Evening complex genes *LUX, ELF3* and *ELF4* were expressed at dusk, whereas *GI*, which plays a role in light-dependent turn-over of the TOC1 protein, was expressed in the late afternoon (Fig. S2). The temporal expression patterns of these clock-associated genes were consistent with those in leaves of *A. thaliana* (1, 37), rice and poplar (38), and the legume soybean (39), suggesting that circadian clock mechanisms are largely conserved between these plant species, and between root nodules and plant leaves.

### Rhythmic coordination of nodule metabolism

Having determined that circadian clock components cycle in nodules, we asked which other aspects of nodule function might be under circadian regulation, through analysis of gene descriptions (obtained from Phytomine in Phytozome) as well as GO term, biological pathway and protein domain enrichment (summarized in Fig. 3G-H). As previously described in *A. thaliana* (2), genes associated with the isoflavonoid pathway were expressed around dawn, including the rate-limiting enzyme phenylalanine ammonia-lyase (PAL). The morning cluster (Cluster 2) was enriched for genes involved in elongation growth, including the transcription factor Phytochrome Interacting Factor 4 (*PIF4*, Medtr3g449770), which promotes auxin biosynthesis in *A. thaliana* (40), eight Walls Are Thin1 (WAT)1-related genes encoding glycoside hydrolases, as well as nodulin genes; all related to cell wall biosynthesis and flavonoid biosynthesis (41) (Fig. 3D). This was consistent with the temporal expression pattern of transcripts associated with cell wall synthesis, both in monocots such as rice and dicots such as poplar (38). The morning cluster also contained 4 genes related to light-harvesting complex I chlorophyll a/b binding proteins, which are associated with plastid function in roots and beneficial plant-microbe interactions (42).

The evening cluster (Cluster 3) contained genes encoding enzymes associated with starch degradation (Fig. 3E). This is consistent with the pattern of starch accumulation and degradation in *A. thaliana*, where starch accumulates during the light period, then is utilized to support growth during the night (3). Three genes related to starch degradation were also expressed in the late-night cluster (Cluster 4, Fig. 3F), indicating continuous starch degradation during the night period, which may be important to provide simple carbon compounds to the rhizobial symbionts.

Late-night and early morning (dawn) processes were largely related to nitrogen metabolism. Both Cluster 4 and 1 contain many genes that are associated with glutamate metabolism, amino acids and nitrate/nitrite transport and Cluster 1 contains glutamine synthetase (Medtr3g065250), which catalyses the first step of nitrogen assimilation (Fig. 3F, Table S4). Furthermore, the ureide biosynthesis pathway was enriched in Cluster 4. Ureides are the main long-distance transport forms of nitrogen from nodules to the shoot and are moved up the xylem vessels to the leaf tissue where then they are used as a nitrogen source (43). Although indeterminate nodule-forming legumes including *M. truncatula* have been classified as amide-type (rather than ureide type), detection of ureide pathway related genes suggests that this part of the metabolism still occurs in these legumes as a response to nitrogen fixation (44). Genes related to phosphate metabolic processes were also enriched in late-night Cluster 4, which may play a critical role to meet the high Pi requirement for nitrogen fixation during the late night (Fig. 3F). Genes related to anion transmembrane transport, over-represented in Cluster 1 (Fig. 3C), may enable movement of compounds formed during the night.

Together, these results indicate that key aspects of nodule function, including starch utlization, nitrogen assimilation and nitrogen transport, occur rhythmically under the control of the circadian clock, and suggest that appropriate temporal coordination of these processes may be important for optimal nodule function.

### Circadian regulation of NCR gene expression in nodules via the Evening Element

We then looked for genes that might play a role to coordinate rhizobial activity with metabolic rhythms in nodules, and found that a large number (45) of NCR transcripts were rhythmic (Fig. 4,Table S5). Rhythmic expression of this group of peptides, known to affect rhizobial activities (25), may be part of the mechanism by which the plant circadian clock impacts on nodulation. The distribution of rhythmic NCRs per cluster showed that there were 2 NCRs in the dawn cluster 1 (0.25%), 5 NCRs in the morning cluster 2 (2%), 24 NCRs in the evening cluster 3 (7.5%) and 14 NCRs in the late-night cluster 4 (1%). These results show an enrichment of NCRs in the evening cluster, compared to a random distribution across all clusters (Fig. 4C, 4.71 over-enriched, *P-*value 1.38e-21).

**Figure 4.**
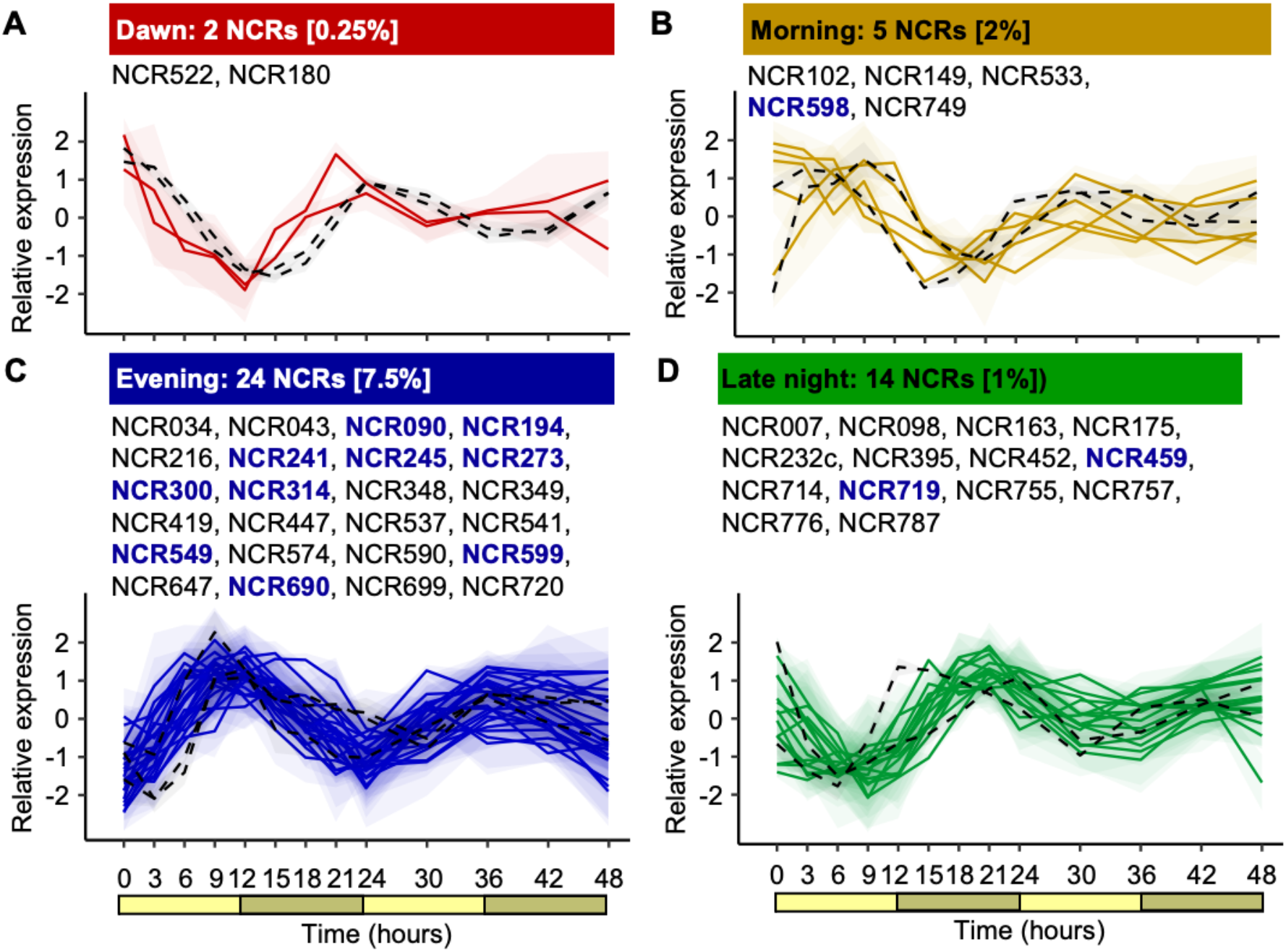
Oscillating expression of NCR genes. (A-D) Presence and expression profiles of NCR (coloured solid lines) genes within each cluster; clock genes within each cluster are shown with black dashed lines, as Fig. 3C-F. The average and range of each group of genes is indicated with lines and a cloud, respectively. NCRs with the EE element in their promoters are indicated with blue bold text. See Table S2 for all data values.

These observations led us to investigate how the clock might control NCR gene expression in nodules. We carried out a *de novo* motif analysis in promoters of NCRs with rhythmic expression (Fig. 5A). This identified 3 over-represented motifs within 500 bp upstream of the translational start sites and within 200 bp of the TATA box (Fig. S3). These 12bp-motifs mapped into longer stretches of conserved sequences identified in a previous study (31).

**Figure 5.**
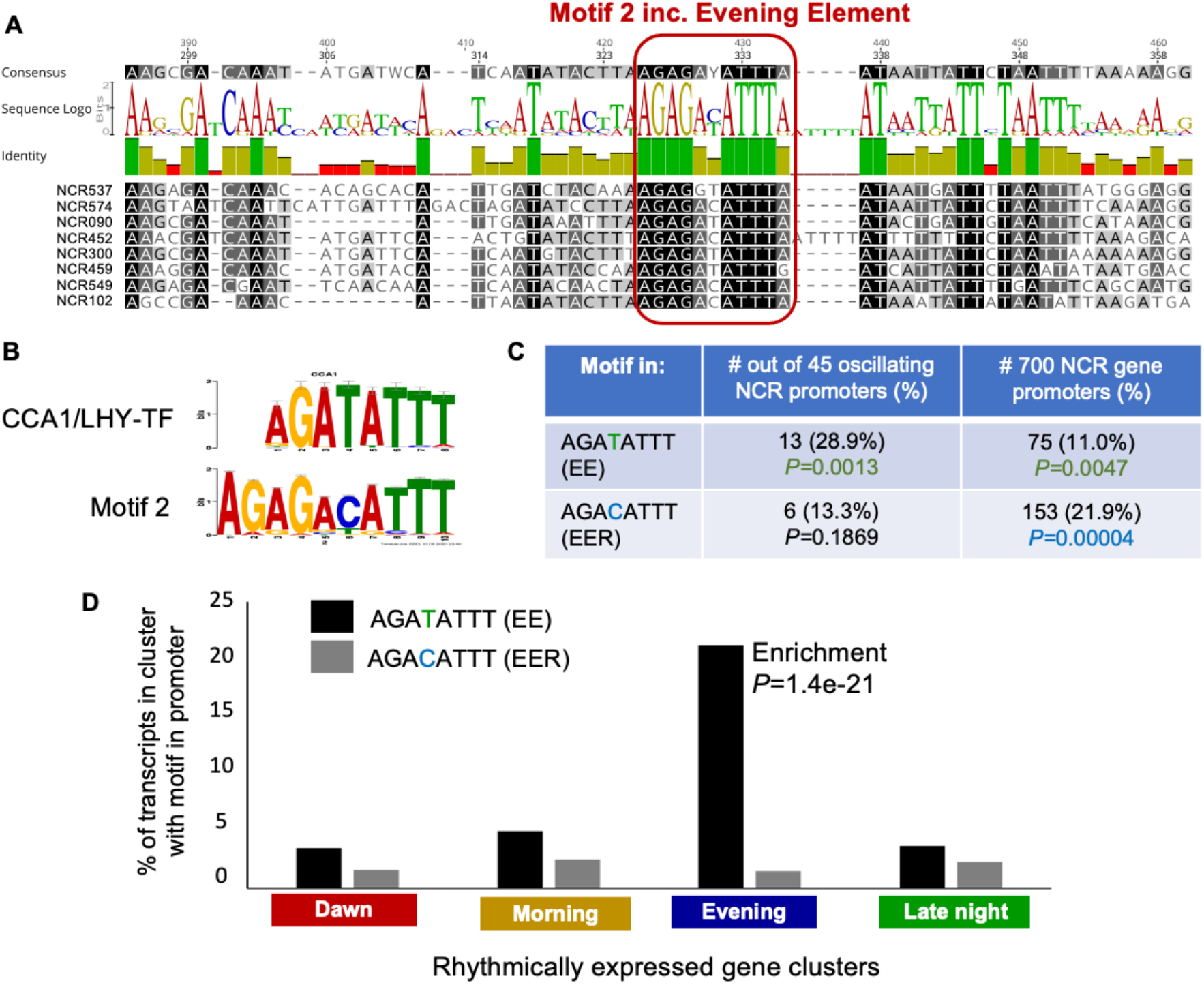
The Evening Element is enriched in the promoters of oscillating NCRs. (A) Visualization of conservation across part of the NCR promoter sequence for an example of eight of the NCRs used for *de novo* motif identification, showing the location of the Evening Element, EE (red box); see Figure S3 for complete promoter alignment for these example NCRs. (B) Motif comparison of the CCA1/LHY binding site with motif 2 (EE/alternative EE, EER). (C) Presence and enrichment (*P* value) of the EE and the EER in the promoters of the 45 oscillating NCRs and the whole family of 700 NCRs; see Table S6 for all motif enrichment statistical analysis. (D) Presence of the EE and the EER in the % of transcript promoters within each cluster; see Table S6 for all motif enrichment statistical analysis.

Motif 2 (AGA[T/C]ATTT, Fig. 5A-B) was closely related to the Evening Element (EE) AGATATTT, which in *A. thaliana* is bound by the circadian clock-associated proteins CCA1 and LHY (2, 34, 35, 45). Consistent with regulation by these dawn-specific transcriptional repressors, more than half of the rhythmic NCRs were enriched in the evening cluster. To obtain further evidence that the EE motif drives rhythmic expression of NCRs, we used the FIMO tool (46) to assess the presence of the motif in the promoters of rhythmic vs non-rhythmic NCRs. This showed that 29% (13) of NCR gene promoters that cycled in our experiment contained the canonical EE motif, AGATATTT, compared to 8% of a random set of non-rhythmic NCR genes (*P*=0.0013 for enrichment) (Fig. 5C, Table S6). We then asked if the EE might also be found in non-rhythmic members of the gene family. Out of the 700 NCR genes, compiled from data from (47) and (48) (Table S5) we identified an EE in the promoters of 75 genes (∼11%), compared to 4% in a randomized set of 700 non-NCR gene promoters (Fig. 5D; *P*=0.0047 for enrichment; Table S6). Almost all promoters that contain the EE had a single occurrence of the motif, with just three promoters having 2 occurrences (Table S5). 63 NCRs contained the EE but did not oscillate in our experiment (Table S5). This is consistent with previous observations that a large proportion of CCA1 and LHY regulatory targets do not exhibit rhythmic expression under any observed condition (34, 35). Expression of these NCRs may cycle at other stages of nodule development or in specific nodule cell types.

The alternative version of the EE motif, AGACATTT (EE-related, or ‘EER’ motif) was only found in 6 rhythmic NCRs (Fig. 5C). To ask whether the EER motif was capable of driving rhythmic gene expression, we searched for this motif in the promoters of the 2,832 cycling transcripts described above (represented by 2,703 genes). Whereas the EE motif was associated with evening-specific gene expression, rhythmic genes containing the EER motif were expressed at all phases of the day-night cycle (Fig. 5D), suggesting that the EER motif might not play a role in circadian regulation of gene expression. On the other hand, the EER motif was found in about 23% of NCR promoters, compared to 2.4% in a control gene set (2.5%, *P*=3.8E-5 for enrichment) suggesting a potential role in nodule-specific gene expression (Fig. 5C). Only 13 NCRs had both EE and EER motifs, supporting the notion that there are distinct functional categories of NCRs (Table S5). Across all NCRs, presence of the EE or of the EER motif was fairly evenly distributed across the phylogeny, suggesting that they did not arise as part of a single lineage-specific expansion event (Fig. S4).

Nodules are comprised of five zones that each have a different role in nodulation, which are the meristem (furthest from the root), the infection zone, the interzone, the nitrogen fixation zone and the senescence zone (closest the root). We asked if the oscillating NCRs are expressed in any of these particular zones by querying data published in a study that determined the expression locations of 521 NCRs out of our list of 701 NCR genes (29). NCR expression is abundant in the interzone (53%), but the interzone was also enriched (*P*=0.039; chi-squared test) for cycling NCRs whose promoter contains an EE, with 77% of these rhythmic NCRs being localized there. Therefore this region of the nodule, site where bacteroids become fully elongated and start to express N-fixation genes along with the expression of some key late nodulins by host cells (49), seems to be particularly enriched for LHY-controlled NCRs. In contrast, NCRs containing the EER motif were not enriched in the interzone (*P*=0.742; chi-squared test). These observations suggest that rhythmic expression of NCRs under the control of the EE may be particularly important at the stage where bacterial symbionts are differentiating into bacteroids.

## Discussion

Many aspects of physiology, metabolism and development exhibit circadian regulation across plants, animals and some microbes. Thus, the clock often influences the outcome of interactions between organisms (8). In plants, the oscillator mechanism of the clock has been investigated at length in shoots (reviewed in (11)). In roots, it is known to be entrained by shoot-derived signals (14). Despite the agricultural importance of the beneficial legume-microbe interaction of nodulation, very little is known about the impact of the plant clock on this nitrogen-fixing symbiosis. In common bean (*Phaseolus vulgaris*), changes in expression levels of clock-associated genes were detected in the early stages of symbiosis, suggesting that the function of the root circadian clock was adjusted in response to infection by rhizobial strains (50). Here we show that loss-of-function of *LHY*, one of the core clock genes, results in reduced nodulation. Plant yield was reduced in *lhy* mutants that were inoculated with rhizobium, but not in un-inoculated plants, suggesting that reduced biomass was caused by disrupted nitrogen fixation. These observations were consistent with a recent report showing that reduced function of *MtLHY* was associated with reduced nodulation (51). While it remains possible that *LHY* impairs nodulation through a development effect, these results suggest that appropriate timing of circadian rhythms in plant nodules may be important for optimal symbiotic interactions with rhizobium.

We show that the temporal pattern of expression of clock-associated genes in *Medicago truncatula* nodules is consistent with that observed in other plant species and in other organs, suggesting that the molecular mechanism of the central oscillator is largely conserved. However, while *LHY* and *CCA1* are closely related and have largely redundant functions in *A. thaliana*, the *M. truncatula* genome contains a single orthologue of these proteins. Whereas loss of function of both genes was required to disrupt free-running rhythmicity in *A. thaliana* (52), loss of function of *MtLHY* led to short period rhythms of leaf movements and gradual arrhythmia in constant light. The LHY binding site, also known as Evening Element or EE, was over-represented in the promoters of *M. truncatula* nodule genes that peaked in expression in the evening. This was consistent with a role for the cognate transcription factor, MtLHY in driving rhythmic gene expression. These findings suggest that MtLHY plays a similar role to its *A. thaliana* orthologue as a core component of the nodule central oscillator.

In order to obtain clues to the mechanisms by which the circadian clock might affect nodulation, we went on to identify rhythmic processes in nodules. About 5% of the genome showed rhythmic expression in constant light, which was much smaller than the fraction reported for whole *A. thaliana* plants, about 15% (1, 2, 53). As no other circadian transcriptome data are available for plant roots or for *M. truncatula*, it is unclear whether this reflects a species difference or a root vs shoot difference. Expression of genes associated with starch degradation was observed in the evening, as previously described in *A. thaliana* (Harmer et al., 2000). This was followed by expression of genes associated with ureide biosynthesis in the late subjective night. Ureides are the main long-distance transport forms of organic nitrogen in legumes, and their production in the late subjective night suggests that nitrogen-fixation in symbiosomes occurs during the night, deriving its energy from starch hydrolysis. In support of this hypothesis, we also find transmembrane amino acid transporters in the late-night/dawn gene clusters.

Genes associated with isoflavonoid biosynthesis peaked around dawn, as previously described in other plants such as Ginkgo (54) and *A. thaliana* ((2)). Isoflavonoids are polycyclic compounds that belong to the wider group of phytoalexins that are synthesized by many plants and many have antimicrobial activities. In legumes, a wide range of isoflavonoid compounds have been described, with the composition mix being different depending on the species (55). Some of these isoflavonoids actually initiate the plant-symbiont molecular dialogue that leads to nodule formation, by inducing the expression of *nod* genes in rhizobia (56). The clock is known to regulate plant defence responses, and plants are typically more resistant to pathogen attacks at dawn (4, 6, 57). Production of flavonoids at dawn may contribute to this gating mechanism, to control entry of microbes into plant roots while attracting rhizobial symbionts. However, flavonoids are thought to have a role beyond initial recruitment of rhizobia, since they are mostly produced in the nodule infection zone, where bacteroids become fully elongated and start to express N-fixation genes (58). In mature nodules, isoflavonoids have been suggested to play a role in maintaining a homogeneous rhizobial population (59). Since expression of genes associated with cell wall biosynthesis peaks in the morning, rhythmic production of flavonoids could act to coordinate nodule cell expansion with bacteroid proliferation at dawn.

Our transcriptomic analysis also revealed the rhythmic expression of a subset of NCRs, with the majority peaking in the evening. This large family of peptides is thought to control bacterial differentiation within the nodule, but there is evidence for functional differentiation of NCRs, as different NCRs can have either pro-symbiotic or anti-symbiotic properties (60, 61) and bacterial elongation and activity in nodules can vary depending on the particular suite of NCRs present in the plant host (47). The observation that a subset of NCRs is expressed rhythmically in nodules suggests a function to synchronize bacterial activity with the rhythms of the plant host and provides further evidence for functional differentiation of this group of peptides. Previous studies of NCR promoters identified long stretches of conserved sequence which included putative regulatory motifs such as an ID1 binding site, an Auxin Response Factor (ARF) binding site, a DOF protein binding site, and MADS transcription factor binding sites (31). Here we show that the EE motif is over-represented within NCR promoters, suggesting direct repression by the MtLHY clock protein in the morning. This explains the temporal expression pattern of the majority of NCRs, peaking in the evening in cluster 3. Some NCRs peaking earlier or later in clusters 2 and 4 that contained EEs in their promoters are also likely to be regulated by MtLHY in combination with other rhythmic transcription factors.

A related motif named EE-related or EER was also identified, which was not associated with expression at specific times of the day but was over-represented in NCR promoters, and thus may be associated with nodule-specific gene expression. This motif was also present within one of the stretches of conserved sequence previously identified in NCR promoters (31). The EE sequence (AGATATTT) lies at the same position within this conserved sequence, but was not uncovered in that previous research, likely due to the EER sequence variant (AGACATTT) being present at a high frequency. The presence of a cytosine in the EER motif is interesting because it could be associated with epigenetic regulation of expression (62).

The coordination of nodule growth with bacterial differentiation and nitrogen fixation in indeterminate legume nodules is a well-orchestrated process. Our results suggest that rhythmic expression of NCR peptides under the control of the plant circadian clock plays a vital role in the establishment of successful symbiotic interactions. Many crops have lost their photoperiodic responses as part of domestication, because this was essential for cultivation at a broad range of latitudes, and in many cases, this happened through disruption of the circadian clock (63). For example, clock components have been modified during the soybean domestication process (reviewed in (64). It is therefore crucial to understand how the clock affects the host-symbiont interaction so we can avoid breeding against the efficiency of the nitrogen fixation process. Moreover, the possibility of modulating specific downstream pathways such as rhythmic NCR peptides may enable optimization of nodulation whilst avoiding undesirable plant clock related side-effects. By identifying a mechanism that links control of plant growth and development with that of its symbiotic partner, our work opens up a new field of investigation for understanding how the rhizobial activity is regulated by the plant.

## Materials and Methods

### Plant materials and growth conditions

*Medicago truncatula* wild-type accession A17 in the Jemalong background was obtained from the IGER seed bank (Aberystwyth, http://www.igergru.ibers.aber.ac.uk). *Tnt1 M. truncatula* mutant lines for *LHY* (Medtr7g118330) in the R108 background were identified from the Noble Research Institute ((https://medicago-mutant.noble.org/mutant/database.php) (65). Lines were identified by querying R108 sequences of the *M. truncatula LHY* coding region plus 200 bp upstream and downstream using a blastn search with default parameter settings (E-value cut-off 10^−6^). *Tnt1* lines were selected based on their E-values and % identity > 95. Lines NF17115 (*lhy-1*) and NF16461 (*lhy-2*) were identified with insertions in the promoter region and fifth exon respectively (Fig. 1B).

Seeds were scarified with concentrated H_2_SO_4_ (∼2-3 mL per 100 seeds) then thoroughly washed 3-5 times with sterile water. Scarified seeds were sterilized by treating with 7% sodium hypochlorite solution for 5 minutes with gentle shaking followed by 7 times repeated washes with sterile water. Seeds were sown on 1.5% phyto-agar plates, sealed using 3M Micropore™ tape, wrapped in foil then left in at 4°C for 72 hrs. Plates were then placed in a Sanyo MLR-352 growth chamber (25°C) for 4 days before seedlings with a radicle length of >2 cm were transferred to FP11 (11×11 cm) pots containing sterilized perlite with a 1-2 cm layer of sterilized vermiculite on top for phenotypic analysis. Pots were placed in a Sanyo 2279 growth cabinet with 12/12 hours light/dark, irradiance of 200 μmol m^−2^ s^−1^ and temperature of 24°C (day) and 21°C (night). Pots were watered 2-3 times a week (when dry) with modified Broughton and Dilworth (1971) nutrient solution (1 mM CaCl_2_, 1 mM KH_2_PO_4_, 75 µM FeNaEDTA, 1mM MgSO_4_, 0.25 mM K_2_SO_4_, 6 µM MnSO_4_, 20 µM H_3_BO_3_, 1 µM ZnSO_4_, 0.5 µM CuSO_4_, 50 nM CoSO_4_, 0.1 µM Na_2_MoO_4_, adjusted to pH 6.5 with KOH). For plant growth for genotyping or seed bulking, germinated seedlings were transferred to FP9 (9×9 cm) pots containing F2 compost and plants grown in a glasshouse compartment at 16/8 hours light/dark, average irradiance of 200 μmol m^−2^ s^−1^ and temperature of 24°C (day) and 21°C (night).

### *lhy* mutant characterization and rhythmic leaf movement assays

For phenotypic analysis, plants were grown under 12L12D for 5 weeks, then removed from pots and photographed before measuring shoot and nodule weights. For rhythmic leaf movement (RLM) assays, plants were grown under 12L12D for 10 days before transferring to constant light for imaging from above using timelapse cameras (Brinno). Opening and closing of the first true leaf was monitored by measuring changes in visible leaf area using ImageJ software. Greyscale images were thresholded and converted to binary with leaves showing white on a dark background. White pixels were then quantified over time in regions of interest using the Integrated Density tool. The experiment was repeated twice, then data from both biological replicates was combined. Baseline detrending was applied to the data and periodicity for the remaining samples was analyzed using FFT-NLLS in BioDare2 (https://biodare2.ed.ac.uk (66)).

### Rhizobial culture preparation and seedling inoculation for timecourse analysis

*Sinorhizobium meliloti* strain WSM1022 was grown on TY/Ca^2+^ plates (5g/L tryptone, 3g/L yeast extract, 6 mM CaCl_2_· 2H_2_O, pH adjusted to 6.8-7.0) at 28°C for 2 days. From the solid culture, *S. meliloti* was spot-inoculated into 10 mL liquid TY/Ca^2+^ medium and grown for ∼24 hours with gentle shaking at 28°C. Rhizobial cells were harvested by centrifugation at 3200g for 10 minutes, washed twice with sterile water, then resuspended in sterile water to an OD_600_=0.05. 250 μL of freshly prepared rhizobial solution was used to inoculate each *M. truncatula* seedling the day after potting by pipetting onto the vermiculite layer in close proximity to the plants.

### Sampling plants for transcriptomic analysis

For RNAseq/qPCR timecourse analysis, after 40 days in 12L12D, pots were transferred to constant light conditions at the same irradiance. At 0hr, and then every 3hr up to 48hr, plants were removed from pots, samples were pooled from 6-7 plants for one biological repetition, immediately flash frozen and stored at −80 °C. Nodules were picked from roots using tweezers, part of each root system without nodules was collected, and leaves were collected as 4-5 trifoliates. Samples for qPCR analysis of *LHY* expression in mutants vs. R108 (Fig. 1B) were taken at 7:30 (morning) and 15:30 (evening) into the light cycle. Experiments were carried out three independent times.

### Genomic DNA extraction and PCR for *Tnt1* line genotyping

*Tnt1 M. truncatula* mutant lines were sterilized, germinated and grown to maturity in a glasshouse compartment. Genomic DNA from a leaf sample from each of the plants were extracted using 5% Chelex suspension column binding and heat treatment (100°C for 5 minutes), then diluted 1/10. Gene-specific primers *lhy-1*Fp (CTCAAAACATGGCGGCTTAC), *lhy-1*Fp (AGTGGCTGAGATTGGTTGTG), *lhy-2*Fp (AATGAACGATTTTAGCAGCGG) and *lhy-2*Rp (TTTGGCCGTATGCAAATGTAG) were designed based on the R108 sequence ∼1000bp away from the FST site for each of the *Tnt1* mutant inserts using Primer3. These gene-specific primers were used in combination with *Tnt1*-specific *Tnt1*-Fg and *Tnt1*-Rg, *Tnt1*-Fg1 (ACAGTGCTACCTCCTCTGGATG) and *Tnt1*-Rg1 (CAGTGAACGAGCAGAACCTGTG) primers for PCR genotyping, using sequences in (67, 68). MyRed Taq DNA polymerase (Bioline) was used in a reaction volume of 20 μl, using touch-down PCR as described in (67).

### RNA extraction, RNAseq and quantitative PCR (qPCR) analysis

Frozen plant tissue samples were finely ground using a mortar and pestle, then around 100mg of each powdered sample was used for total RNA extraction then gDNA removal, using the Monarch® Total RNA Miniprep Kit. The quantity (>100 ng/μl) and quality (RNA integrity > 8.5) of RNA was determined using a Bioanalyzer 2100 RNA 6000 Pico Total RNA Kit (Agilent Technologies). Samples containing >5 μg of RNA in total were used for RNAseq. mRNA library preparation, quality assessment and sequencing (150bp, unstranded, paired-end) were carried out by Novogene; mRNA libraries were prepared following the Illumina TruSeqTM RNA library preparation protocol, after rRNA had been removed using the Ribo-Zero kit.

For qPCR analysis, cDNA was prepared using the ProtoScript II First Strand cDNA Synthesis Kit from New England Biolabs (UK) Ltd. qPCR was performed with 20 µl reaction volumes using SYBR Green JumpStart Taq ReadyMix (Sigma-Aldrich) and 40 two-step amplification cycles (95°C and 60°C for 30s and 60s, respectively) in a 96-well Agilent Mx3005P real time PCR machine. Primer pairs designed based on the borders of the 2^nd^ and 3^rd^ exons (Fp: CACAAAACAAAGAGAACGATGG, Rp: ATGGCTCCTGATTTGCACAG) were, used for the quantification of *LHY* expression, normalized against the reference gene Mtβ-Tubulin Medtr7g089120 (Fp: TTTGCTCCTCTTACATCCCGTG, Rp: GCAGCACACATCATGTTTTTGG). Data was analyzed using the ΔC_t_ method, a derivation of the ΔΔC_t_ method (69).

### Statistical analysis of transcriptomic levels

Raw sequence data in the form of a pair of .fq.gz files with sequencing depth of at least 20 million reads per sample were processed using tools on the Galaxy EU server (usegalaxy.eu). First, the quality of raw sequencing data was analyzed by FastQC (70). Replicate 2 for 21hr and replicate 3 for 15hr were found to have poor quality data and were removed from analysis. Contaminating adapter sequences and poor-quality sequences were removed by Trimmomatic v36.4 (71) with the following settings: slidingwindow:4:20 and minlen:40 and using phred33 quality scores. Next, these clean, trimmed and paired reads were used to generate raw transcript read-counts and transcripts per million (TPM) normalized read-counts using Salmon quant v0.14.1 (72) with *M. truncatula* reference transcript sequences (Mt4.0v1) downloaded from the Phytozome database (phytozome.jgi.doe.gov). Expression data for a total of 61,510 transcripts was generated. Read counts were further normalized as log2 transcripts per million (logTPM). To identify the diurnally oscillating transcripts, logTPM expression data were analyzed using the R package MetaCycle v1.2.0 (73) with the following settings: minper: 20, maxper: 28, cycMethod: LS (Lomb-Scargle).

Hierarchical clustering using the total within-cluster sum of square (elbow method) was performed in R using 1-Pearson’s correlation coefficient as a dissimilarity distance measure between normalized (mean centred and scaled by SD), oscillating genes. Enrichment analysis for processes was performed with Bonferroni method of correction (*P*-values<0.05) detailed in Table S4.

### Promoter motif presence and structure analysis

Promoter sequences of *M. truncatula* (Mt4.0v1) genes were retrieved from the *M. truncatula* genome database (http://www.medicagogenome.org). Promoter motif analysis was carried using the MEME suite (74) and *de novo* motif discovery runs were performed on either strand of unaligned 500 bp upstream sequence with motif width of 12 bp. We subsequently also queried 200bp and 1000bp of each NCR promoter, finding that motifs were clustered within the 500bp region; this location is consistent with findings from (31). Conserved motifs were selected based on bit size (range from 0-2), positional bias (*P-*value<0.05) and with an E-value<0.001 (Fig. S3).

Position weight matrix (PWM) scanning for known motifs was performed using Find Individual Motif Occurrence (FIMO) (46) to determine the number of matches in each promoter region. FIMO was used to search for the occurrence of MEME-generated motifs against these backgrounds on 500 bp promoter upstream sequences with a *P*-value<1E-4 (46), then genes with at least one motif site in their promoter were assessed for enrichment. For analysis of the whole NCR gene family, 700 NCRs (compiled from data from (47) and (48), Table S5) against 3 sets of 700 randomly sampled non-NCRs from the remainder of the *M. truncatula* genome were used for enrichment analysis. The Shapiro-Wilk test was used to determine enrichment data normality and an F-test was used to determine data variance. A two sample Welch t-test and pairwise Wilcox test was then used to calculate *P*-values in R. Similarly, enrichment for the occurrence of the EE in the 45 cycling NCR genes was compared to: (i) 45 expressed but non-cycling NCRs (ii) 45 expressed and cycling non-NCRs and (iii) 45 non-expressed and non-cycling NCRs. FIMO scan for AGATATT enrichment was also performed on upstream regions of (i) all 2,832 cycling transcripts (ii) randomly sampled sets of 2,832 non-cycling genes. For all FIMO enrichment analyses, MEME was used to generate motifs with a size of 10 bp nucleotides and random samples were generated in R.

### Multiple sequence alignment of NCR promoters

Promoter sequences were aligned using MAFFT (Multiple Alignment using Fast Fourier Transform) tools at EBI (European Bioinformatics Institute) (75). The promoters of eight cycling NCR genes were aligned using a Hidden Markov Model (HMM), selected based on the occurrence of all the three motif sites. For optimal alignment and representation of motif conservation, we ran algorithms with a parameter setting of gap lowest open penalty 1 allowed in MAFFT, gap extension 0.5 and an iteration of 100 runs; due to the diverse nature of the NCR sequences (76), gaps were required to generate the best local and global alignment (77). The aligned sequences were then visualized in Genious version 11.0.2 (https://www.geneious.com).

### Transcription factor search and ortholog identification

Tomtom (78), was used to search for PWM query motifs against the *A. thaliana* PBM db (79), DAP motifs (O’ Malley 2016) and JASPAR plants 2018 databases of known transcription factor binding sites. For othologous gene identification in *M. truncatula*, we used two methods. Firstly, reciprocal BLASTp was performed with the NCBI blasp suite, them alignment score, percentage query coverage and expect value were determined for forward and reciprocal queries (80). Secondly, a Smith-Watermann (SW) alignment homologue search was carried out using the Phytozome v12.1.6 database. Different homologs of same clock gene are marked as a/b/c, based on their level of similarity to *A. thaliana* ortholog, with a being the highest. Although we did not find orthologs for *PRR9* and *PRR3*, it should be noted that *M. truncatula* homologs listed under *PRR7/5* also share similarity with *AtPRR9/3*.

### Promoter phylogenetic tree reconstruction

The DNA sequences of 700 NCRs with their upstream region were aligned using MPI-based MAFFT version 7.3 https://mafft.cbrc.jp/alignment/server/index.html for large sequences. Except for three NCR genes (Medtr0538s0010, Medtr1886s0010, Medtr0753s0010) in scaffolds that have an upstream region <500 bp, all other NCR promoters were of 500 bp upstream length. Maximum-likelihood (ML) analyses and search for the best-scoring tree were performed using RAxML version 8.2.10 with rapid bootstrapping of 100 replica runs. The substitution model of Generalised Time Reversal (GTR) and the Gamma model of rate heterogeneity was used. The best resulting ML tree for DNA alignments was used for visualization of the phylogenetic tree on FigTree version 1.4.4. The presence of the EE (AGATATTT), AGACATTT or both AGAC/TATTT in the promoter for each NCRs were then highlighted manually with colours on FigTree.

### CCA1/LHY phylogenetic tree reconstruction

Clock gene homologues were initially identified via reciprocal BLASTp was performed with the NCBI blasp suite of *A. thaliana* clock protein sequences against sequences in the National Center for Biotechnology Information (NCBI). Hits were sorted primarily by maximum bitscore score followed by E value. In cases where there were multiple high scoring hits, multiple putative orthologues were identified. Homology was also assessed using the Smith-Watermann (SW) alignment homologue search with the Phytozome v12.1.6 database. Evolutionary history was inferred using the Maximum Likelihood method and JTT matrix-based model (81). Initial trees for the heuristic search were obtained by applying Neighbor-Join and BioNJ algorithms to a matrix of pairwise distances estimated using the JTT model. Trees with superior log likelihoods are shown. Trees were visualised using the interactive tree of life v4 (82).

## Author Contributions

M.LG., B.L., S.O. and I.A.C. designed research; M.A, P.R., B.L., R.B-P., B.R., N.A., L.B., E.P., J.B., M.R., E.F. and C.R-G. performed research; J.W and K.S.M developed materials; M.A, P.R., A.P., B.L., S.O., I.A.C and M.L.G analysed data; and M.A, P.R., B.L, I.A.C. and M.L.G. wrote the paper.

## Data sharing

All raw RNAseq data has been deposited in the NCBI SRA database (PRJNA634620)

## Acknowledgments

This study was supported by grants from the Biotechnology and Biological Sciences Research Council (BBSRC; BB/P002145/1 to M.L.G and BB/T015357/1 to M.L.G and I.A.C), PhD studentships from BBSRC through the Midlands Integrative Bioscience Training Partnership to M.A. and B.R, from Warwick Chancellor’s International Scholarship Scheme to R.B. and from the International Commonwealth Commission to P.R. The *Medicago truncatula* plants utilised in this research project, which are jointly owned by the Centre National De La Recherche Scientifique, were obtained from Noble Research Institute, LLC and were created through research funded, in part, by grants from the National Science Foundation (DBI-0703285 and IOS-1127155).

---

**Figure S1.**
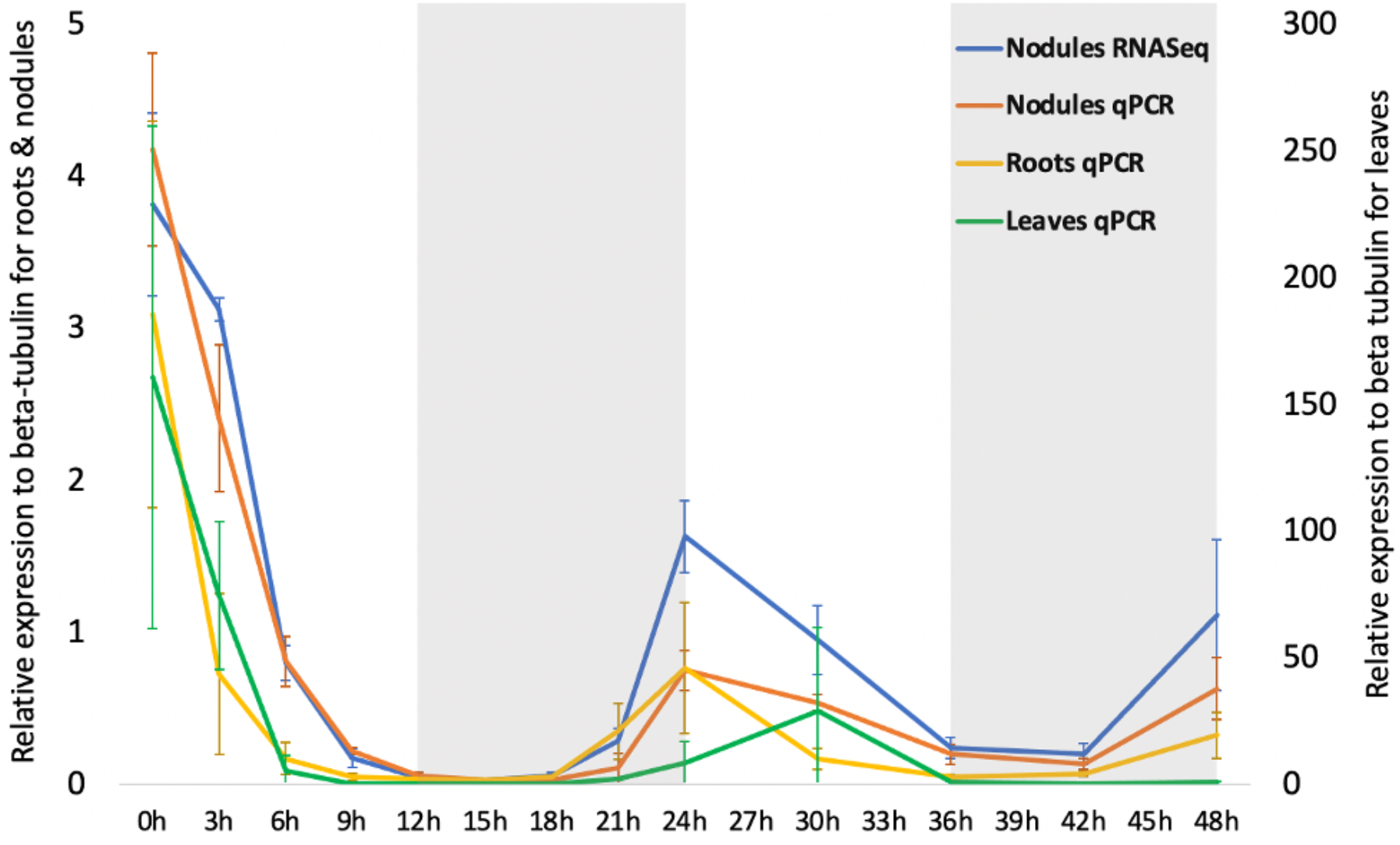
*LHY* is rhythmically expressed in nodules. *LHY* expression exhibits free-running rhythmicity with a dawn peak of expression, consistent with the presence of a functional clock in nodules. The overall expression of *LHY* is highest in leaves, as expected, but the same rhythm is visible in leaves (green, right-hand y-axis), roots (yellow, left-hand y-axis) and nodules (orange, left-hand y-axis); all measured using qPCR across the timecourse. Expression measurement in nodules using qPCR (orange) corroborates the measurement using RNAseq (blue). White and grey background represents presumptive day and night; error bars indicate standard deviation of the mean (n=3 biological replicates); see Table S3 for all values.

**Figure S2.**
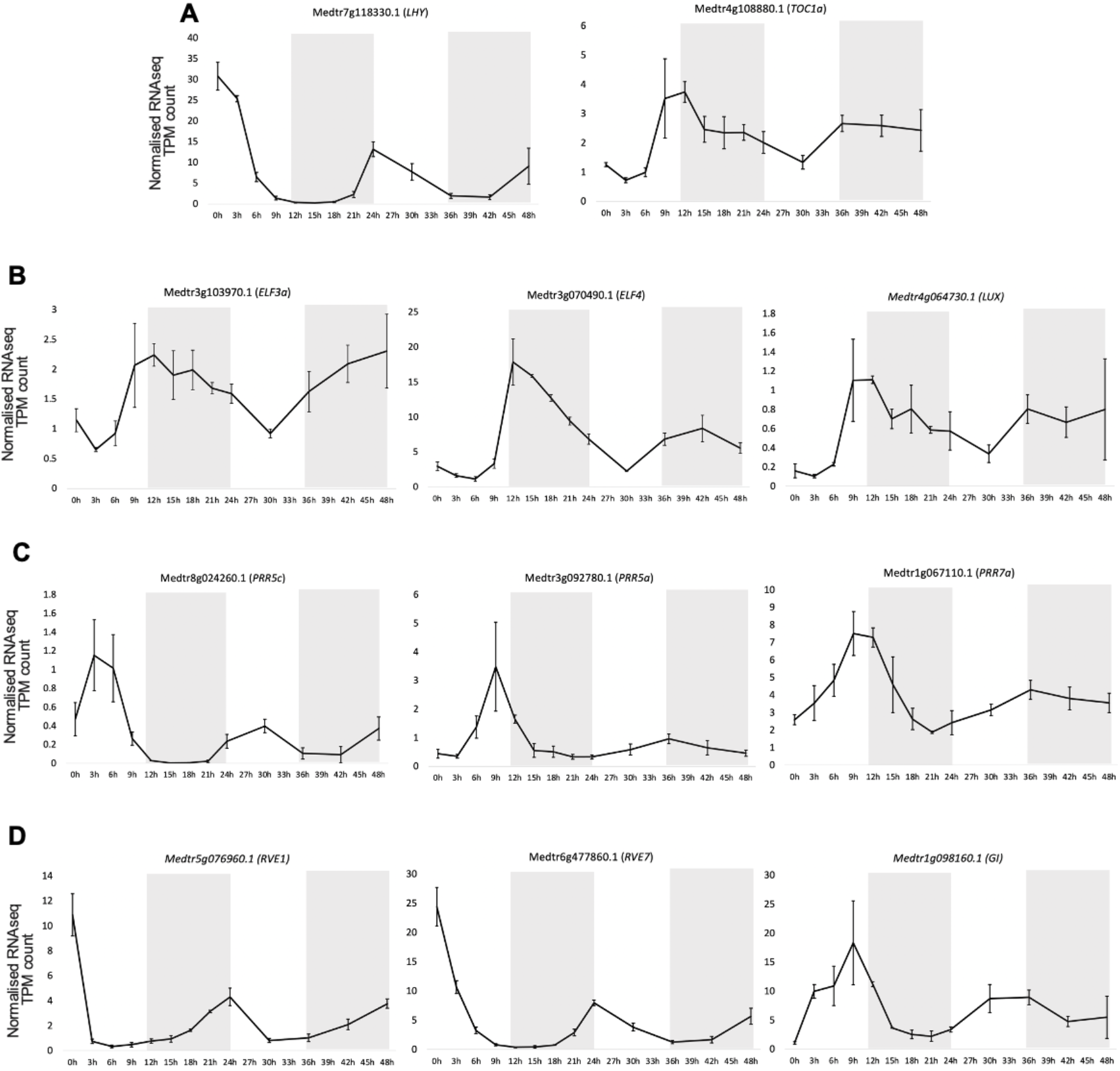
Oscillation of circadian clock genes over nodule timecourse. (A) The core oscillator: LHY/CCA1 and TOC1 form the core oscillatory loop via a negative feedback mechanism. (B) The Evening Complex (EC) is formed of three proteins, ELF3, ELF4 and LUX, whose expression peaks in the evening; (C) Pseudo-response regulators (PRRs) are homologous to TOC1 and expressed mainly during the day to form additional loops. (D) RVEs and Gigantea (GI) synchronize the central clock with output pathways. White and grey background represents presumptive day and night (plants were transferred from 12/12 hours dark/light to constant light before sampling); error bars indicate standard deviation of the mean (n=3 biological replicates, except for 15h and 21h where n=2); all graphs have the same y-axis (normalised RNAseq transcripts per million (TPM) count). See Table S2-S3 for all data values.

**Figure S3:**
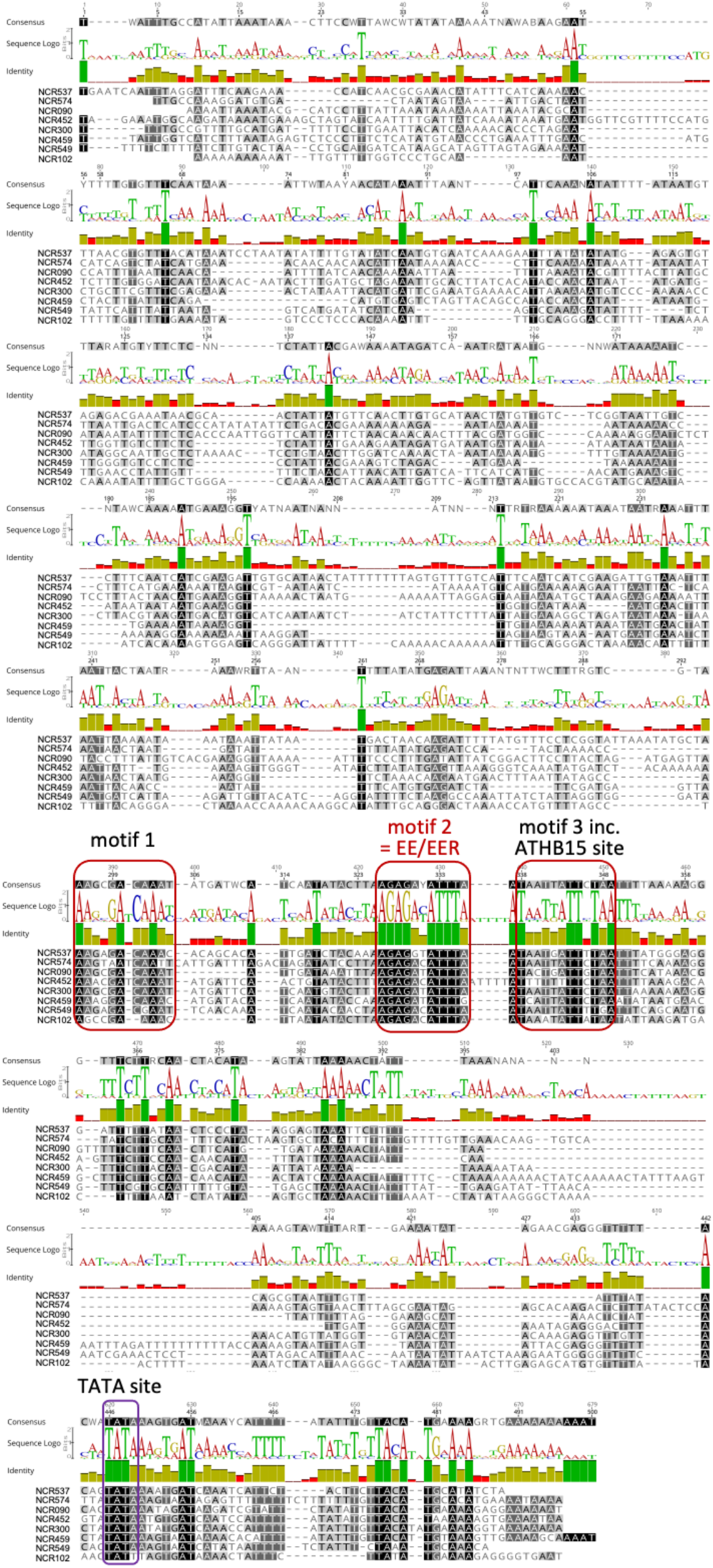
Promoter landscape of the NCRs. Multiple Sequence alignment (MSA) visualization of the conserved motifs in eight NCR promoters that best represents the four conserved regions (red boxes) using Genious 1102. The visualization was performed after aligning 500bp upstream promoters using the EBI tool MAFFT aligned with open gap and gap extension penalty due to the diverse nature of the NCR sequences (see Methods). Conserved motifs were selected based on bits size (range from 0-2), positional bias (*P*-value<0.05) and with an E-value< 0.001. The evening element (EE) was found within the conserved region that we labelled ‘motif 2’, there is some homology between the binding sites of ATHB15/ATHB16 and motif 3, and with the binding site of AHL20 and motif 4. *de novo* identified motif 1 has no known transcriptional regulator. The TATA binding box is indicated within a purple box.

**Figure S4.**
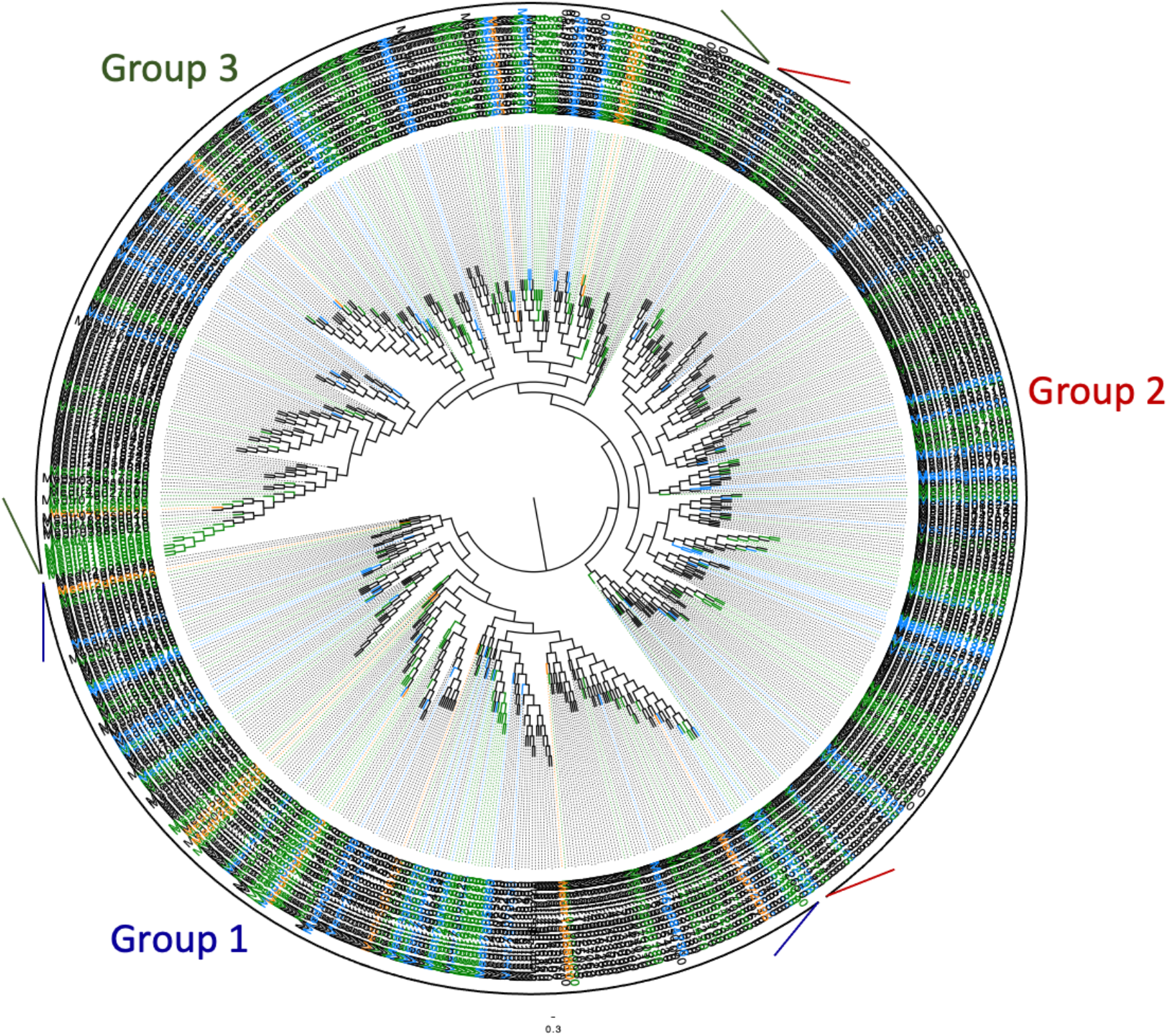
Phylogenetic tree of NCR promoter sequences shows high levels of similarity across the family. Polar visualisation of the promoters of 700 NCRs that all have promoter regions of 500bp, excepting three scaffolds (Medtr0538s0010, Medtr1886s0010, Medtr0753s0010). Blue colour denotes NCRs with the evening element, EE (AGATATTT) in their promoter region, green colour with the EER motif AGACATTT and orange colour denotes NCRs with both the EE and EER motif. The NCR promoters can be broadly divided into three groups (denoted 1,2,3 from the root outwards) in which there are NCRs with the EE or the EER. Promoters with both the EE and the EER (orange) are only present in groups 1 and 3. The tree represents DNA maximum-likelihood phylogeny of NCR promoters with 100 bootstrapping and is rooted in the middle. Scale bar of 0.3 represents branch length, measured as the rate of nucleotide substitution per site.

**Table S1. Analysis of the *lhy-1* and *lhy-2* mutants**. (A) qPCR comparison of *LHY* expression (relative fold change to β-tubulin) in the *lhy-1* and *lhy-2* mutants compared to WT R108. (B) Visible leaf area and period length for *lhy-1* and *lhy-2* mutants compared to WT R108. (C) Nodule fresh weight and plant shoot dry weight for *lhy-1* and *lhy-2* mutants compared to WT R108, in mock-inoculated or *S. meliloti* WSM1022-inoculated conditions. (D) Expression of *LHY, PRR5a* and *ELF4a* transcripts in WT A17 RNAseq data. Rhythmicity was assessed using MetaCycle, and PhaseShift, PhaseShiftHeight, PeakIndex, PeakSPD, Period and *P*-value for oscillation are given. (E) List of geneIDs for LHY/CCA1 homologues, as shown in Fig. 1A.

**Table S2. *Medicago truncatula* A17 transcript IDs detected in 40 day-old A17 nodules and information on their oscillating behavior**. *Medicago truncatula* transcript IDs identified in RNAseq table and information on oscillation pattern (Yes or No), oscillating cluster number (1-4), gene description (from Phytomine), presence of motifs (EE motif=AAGATATTT and EER motif=AAGACATTT) tested if genes are oscillating, oscillating *P*-values (LS_*P-*value and LS_BH.Q *P*-value-FDR correction), oscillating period (LS_period), phase (LS_adjphase), and amplitude (LS_amplitude), log2-transformed RNAseq expression values (TPM) for each transcript, timepoint and biological repeat (columns M-AW), average log2 RNAseq expression values per timepoint (columns AX-BJ), average log2 gene expression value for each gene (BK) and average log2 RNAseq expression values per timepoint compared to the mean for each gene (columns BL-BX).

**Table S3. Expression of clock gene transcripts detected in 40 day-old A17 nodules and information on their oscillating behavior**. (A) *Medicago truncatula* proteins identified with homology to *Arabidopsis thaliana* clock genes returned from reciprocal BLASTp and Smith-Watermann (SW) alignment homologue searches. Reciprocal BLASTp was performed with the NCBI blasp suite, and alignment score, percentage query coverage and expect value are reported for forward and reciprocal queries. Smith-Watermann alignments were retrieved from the Phytozome v12.1.6 database, SW alignment scores, forward and reciprocal similarity and the incidence of a reciprocal best hit are reported. In cases where a reciprocal high degree of similarity was identified, multiple potential homologues are identified. For each transcript, information on oscillation pattern (Yes or No), oscillating cluster number (1-4), oscillating *P-*values (LS_pvalue and LS_BH.Q *P-*value-FDR correction), oscillating period (LS_period), phase (LS_adjphase), and amplitude (LS_amplitude), average and standard deviation (SD) of log2 RNAseq expression values (TPM) per timepoint. (B-C) *LHY* expression measured in leaves, roots and nodules; C are log2 values of B.

**Table S4. GO term, pathway and protein domain term enrichment in oscillating gene clusters**. (A) Terms, *P-*value of enrichment with Bonferroni testing and genes associated with each term for dawn cluster 1 (A), morning cluster 2 (B), evening cluster 3 (C) and late-night cluster 4 (D).

**Table S5. Genes annotated as NCRs in Medicago truncatula A17 and information on their expression levels, oscillating behaviour and presence of motifs**. *Medicago truncatula* transcript IDs identified in RNAseq table, NCR number (where annotated), gene description (from Phytomine), peptide sequence and length, and information on oscillation pattern (Yes (yellow shading) or No), cycling *P*-value (from Metacycle) and oscillating cluster number (1-4) (A-H). Information on presence of motifs (EE motif=AGATATTT and EER motif=AGACATTT) together with matched sequence, location and *P*-values (I-AD). Average log2-transformed RNAseq expression values (TPM) for each timepoint (columns AE-AQ), average log2 gene expression value for each gene (AR) and average log2 RNAseq expression values per timepoint compared to the mean for each gene (columns AS-BE).

**Table S6. Motif analysis in NCRs and cycling genes**. (A) Motifs found in the promoters of rhythmic NCRs. (B) Presence of the EE and the EER in the promoters of the 45 cycling NCRs compared to randomized sets of 45 other promoters (as Fig.3G). (C) Presence of the EE and the EER in the promoters of the 700 NCRs compared to randomized sets of 700 other promoters (as Fig.1E). (D) Presence of the EE and the EER in the promoters of the 2,832 cycling NCRs (as Fig.3G). (E) Presence of the EE and the EER within the four clusters of cycling gene promoters (as Fig.3H). For enrichment analysis see Methods; *P*-values in green are <0.05.

## References

1. T.P. Michael, T.C. Mockler, G. Breton, C. McEntee, A. Byer, J.D. Trout, S.P. Hazen, R. Shen, H.D. Priest, C.M. Sullivan, S.A. Givan, M. Yanovsky, F. Hong, S.A. Kay., J. Chory, Network discovery pipeline elucidates conserved time-of-day-specific cis-regulatory modules. PLoS Genet. 4, e14 (2008).

2. S.L. Harmer, J.B. Hogenesch, M. Straume, H.S. Chang, B. Han, T. Zhu, X. Wang, J.A. Kreps., S.A. Kay, Orchestrated transcription of key pathways in Arabidopsis by the circadian clock. Science 290, 2110–2113 (2000).

3. A. Graf, A. Schlereth, M. Stitt., A.M. Smith, Circadian control of carbohydrate availability for growth in Arabidopsis plants at night. Proc. Natl. Acad. Sci. U.S.A. 107, 9458–9463 (2010).

4. V. Bhardwaj, S. Meier, L.N. Petersen, R.A. Ingle., L.C. Roden, Defence responses of Arabidopsis thaliana to infection by Pseudomonas syringae are regulated by the circadian clock. PLoS One 6, e26968 (2011).

5. R.A. Ingle, C. Stoker, W. Stone, N. Adams, R. Smith, M. Grant, I. Carre, L.C. Roden., K.J. Denby, Jasmonate signalling drives time-of-day differences in susceptibility of Arabidopsis to the fungal pathogen Botrytis cinerea. Plant J. 84, 937–948 (2015).

6. H. Lu, C.R. McClung., C. Zhang, Tick Tock: Circadian regulation of plant innate immunity. Annu. Rev. Phytopathol. 55, 287–311 (2017).

7. C. Zhang, Q. Xie, R.G. Anderson, G. Ng, N.C. Seitz, T. Peterson, C.R. McClung, J.M. McDowell, D. Kong, J.M. Kwak., H. Lu, Crosstalk between the circadian clock and innate immunity in Arabidopsis. PLoS. Pathog. 9, e1003370 (2013).

8. M.J. de Leone, C.E. Hernando, A. Romanowski, D.A. Careno, A.F. Soverna, H. Sun, N.G. Bologna, M. Vazquez, K. Schneeberger., M.J. Yanovsky, bacterial infection disrupts clock gene expression to attenuate immune responses. Curr. Biol. 30, 1740–1747 e1746 (2020).

9. C.J. Hubbard, M.T. Brock, L.T. van Diepen, L. Maignien, B.E. Ewers., C. Weinig, The plant circadian clock influences rhizosphere community structure and function. ISME J. 12, 400– 410 (2018).

10. C. Staley, A.P. Ferrieri, M.M. Tfaily, Y. Cui, R.K. Chu, P. Wang, J.B. Shaw, C.K. Ansong, H. Brewer, A.D. Norbeck, M. Markillie, F. do Amaral, T. Tuleski, T. Pellizzaro, B. Agtuca, R. Ferrieri, S.G. Tringe, L. Pasa-Tolic, G. Stacey., M.J. Sadowsky, Diurnal cycling of rhizosphere bacterial communities is associated with shifts in carbon metabolism. Microbiome 5, 65 (2017).

11. C.R. McClung, The plant circadian oscillator. Biology (Basel) 8, 14 (2019).

12. A.B. James, J.A. Monreal, G.A. Nimmo, C.L. Kelly, P. Herzyk, G.I. Jenkins., H.G. Nimmo, The circadian clock in Arabidopsis roots is a simplified slave version of the clock in shoots. Science 322, 1832–1835 (2008).

13. S. Bordage, S. Sullivan, J. Laird, A.J. Millar., H.G. Nimmo, Organ specificity in the plant circadian system is explained by different light inputs to the shoot and root clocks. New Phytol. 212, 136–149 (2016).

14. N. Takahashi, Y. Hirata, K. Aihara., P. Mas, A hierarchical multi-oscillator network orchestrates the Arabidopsis circadian system. Cell 163, 148–159 (2015).

15. C. Bendix, C.M. Marshall., F.G. Harmon, Circadian clock genes universally control key agricultural traits. Mol. Plant 8, 1135–1152 (2015).

16. M. Li, L. Cao, M. Mwimba, Y. Zhou, L. Li, M. Zhou, P.S. Schnable, J.A. O’Rourke, X. Dong., W. Wang, Comprehensive mapping of abiotic stress inputs into the soybean circadian clock. Proc. Natl. Acad. Sci. U.S.A. 116, 23840–23849 (2019).

17. Y. Wang, L. Yuan, T. Su, Q. Wang, Y. Gao, S. Zhang, Q. Jia, G. Yu, Y. Fu, Q. Cheng, B. Liu, F. Kong, X. Zhang, C.P. Song, X. Xu., Q. Xie, Light-and temperature-entrainable circadian clock in soybean development. Plant Cell Environ. 43, 637–648 (2020).

18. J. Weiss, M.I. Terry, M. Martos-Fuentes, L. Letourneux, V. Ruiz-Hernandez, J.A. Fernandez., M. Egea-Cortines, Diel pattern of circadian clock and storage protein gene expression in leaves and during seed filling in cowpea (Vigna unguiculata). BMC Plant Biol. 18, 33 (2018).

19. J.L. Weller., R. Ortega, Genetic control of flowering time in legumes. Front. Plant Sci. 6, 207 (2015).

20. V. Hecht, F. Foucher, C. Ferrandiz, R. Macknight, C. Navarro, J. Morin, M.E. Vardy, N. Ellis, J.P. Beltran, C. Rameau., J.L. Weller, Conservation of Arabidopsis flowering genes in model legumes. Plant Physiol. 137, 1420–1434 (2005).

21. S.B. Preuss, R. Meister, Q. Xu, C.P. Urwin, F.A. Tripodi, S.E. Screen, V.S. Anil, S. Zhu, J.A. Morrell, G. Liu, O.J. Ratcliffe, T.L. Reuber, R. Khanna, B.S. Goldman, E. Bell, T.E. Ziegler, A.L. McClerren, T.G. Ruff., M.E. Petracek, Expression of the Arabidopsis thaliana BBX32 gene in soybean increases grain yield. PLoS One 7, e30717 (2012).

22. G. Maroti, E. Kondorosi, Nitrogen-fixing Rhizobium-legume symbiosis: are polyploidy and host peptide-governed symbiont differentiation general principles of endosymbiosis? Front. Microbiol. 5, 326 (2014).

23. J.I. Sprent, E.K. James, Legume evolution: where do nodules and mycorrhizas fit in? Plant Physiol. 144, 575–581 (2007).

24. N.D. Young, F. Debelle, G.E. Oldroyd, R. Geurts, et al., The Medicago genome provides insight into the evolution of rhizobial symbioses. Nature 480, 520–524 (2011).

25. P. Roy, M. Achom, H. Wilkinson, B. Lagunas., M.L. Gifford, Symbiotic efficiency enabled by diversification … from 7 to over 700 Nodule-specific Cysteine-Rich peptides. Genes 11, 348 (2020).

26. P. Czernic, D. Gully, F. Cartieaux, L. Moulin, I. Guefrachi, D. Patrel, O. Pierre, J. Fardoux, C. Chaintreuil, P. Nguyen, F. Gressent, C. Da Silva, J. Poulain, P. Wincker, V. Rofidal, S. Hem, Q. Barriere, J.F. Arrighi, P. Mergaert., E. Giraud, Convergent evolution of endosymbiont differentiation in Dalbergioid and Inverted repeat-lacking clade legumes mediated by nodule-specific cysteine-rich peptides. Plant Physiol. 169, 1254–1265 (2015).

27. S. Nallu, K.A. Silverstein, P. Zhou, N.D. Young., K.A. Vandenbosch, Patterns of divergence of a large family of nodule cysteine-rich peptides in accessions of Medicago truncatula. Plant J. 78, 697–705 (2014).

28. I. Guefrachi, M. Nagymihaly, C.I. Pislariu, W. Van de Velde, P. Ratet, M. Mars, M.K. Udvardi, E. Kondorosi, P. Mergaert., B. Alunni, Extreme specificity of NCR gene expression in Medicago truncatula. BMC Genomics 15, 712 (2014).

29. B. Roux, N. Rodde, M.F. Jardinaud, T. Timmers, L. Sauviac, L. Cottret, S. Carrere, E. Sallet, E. Courcelle, S. Moreau, F. Debelle, D. Capela, F. de Carvalho-Niebel, J. Gouzy, C. Bruand., P. Gamas, An integrated analysis of plant and bacterial gene expression in symbiotic root nodules using laser-capture microdissection coupled to RNA sequencing. Plant J. 77, 817–837 (2014).

30. P. Mergaert, K. Nikovics, Z. Kelemen, N. Maunoury, D. Vaubert, A. Kondorosi., E. Kondorosi, A novel family in Medicago truncatula consisting of more than 300 nodule-specific genes coding for small, secreted polypeptides with conserved cysteine motifs. Plant Physiol. 132, 161–173 (2003).

31. S. Nallu, K.A. Silverstein, D.A. Samac, B. Bucciarelli, C.P. Vance., K.A. VandenBosch, Regulatory patterns of a large family of defensin-like genes expressed in nodules of Medicago truncatula. PLoS One 8, e60355 (2013).

32. E. Kondorosi, P. Mergaert., A. Kereszt, A paradigm for endosymbiotic life: cell differentiation of Rhizobium bacteria provoked by host plant factors. Annu. Rev. Microbiol. 67, 611–628 (2013).

33. B. Lagunas, M. Achom, R. Bonyadi-Pour, A.J. Pardal, B.L. Richmond, C. Sergaki, S. Vazquez, P. Schafer, S. Ott, J. Hammond., M.L. Gifford, Regulation of resource partitioning coordinates nitrogen and rhizobia responses and autoregulation of nodulation in Medicago truncatula. Mol. Plant 12, 833–846 (2019).

34. S. Adams, J. Grundy, S.R. Veflingstad, N.P. Dyer, M.A. Hannah, S. Ott., I.A. Carre, Circadian control of abscisic acid biosynthesis and signalling pathways revealed by genome-wide analysis of LHY binding targets. New Phytol. 220, 893–907 (2018).

35. D.H. Nagel, C.J. Doherty, J.L. Pruneda-Paz, R.J. Schmitz, J.R. Ecker., S.A. Kay, Genome-wide identification of CCA1 targets uncovers an expanded clock network in Arabidopsis. Proc. Natl. Acad. Sci. U.S.A. 112, E4802–4810 (2015).

36. G. Wu, R.C. Anafi, M.E. Hughes, K. Kornacker., J.B. Hogenesch, MetaCycle: an integrated R package to evaluate periodicity in large scale data. Bioinformatics 32, 3351– 3353 (2016).

37. A. Pokhilko, A.P. Fernandez, K.D. Edwards, M.M. Southern, K.J. Halliday., A.J. Millar, The clock gene circuit in Arabidopsis includes a repressilator with additional feedback loops. Mol. Syst. Biol. 8, 574 (2012).

38. S.A. Filichkin, G. Breton, H.D. Priest, P. Dharmawardhana, P. Jaiswal, S.E. Fox, T.P. Michael, J. Chory, S.A. Kay., T.C. Mockler, Global profiling of rice and poplar transcriptomes highlights key conserved circadian-controlled pathways and cis-regulatory modules. PLoS One 6, e16907 (2011).

39. J. Marcolino-Gomes, F.A. Rodrigues, R. Fuganti-Pagliarini, C. Bendix, T.J. Nakayama, B. Celaya, H.B. Molinari, M.C. de Oliveira, F.G. Harmon., A. Nepomuceno, Diurnal oscillations of soybean circadian clock and drought responsive genes. PLoS One 9, e86402 (2014).

40. K.A. Franklin, S.H. Lee, D. Patel, S.V. Kumar, A.K. Spartz, C. Gu, S. Ye, P. Yu, G. Breen, J.D. Cohen, P.A. Wigge., W.M. Gray, Phytochrome-interacting factor 4 (PIF4) regulates auxin biosynthesis at high temperature. Proc. Natl. Acad. Sci. U.S.A. 108, 20231–20235 (2011).

41. P. Ranocha, N. Denance, R. Vanholme, A. Freydier, Y. Martinez, L. Hoffmann, L. Kohler, C. Pouzet, J.P. Renou, B. Sundberg, W. Boerjan., D. Goffner, Walls are thin 1 (WAT1), an Arabidopsis homolog of Medicago truncatula NODULIN21, is a tonoplast-localized protein required for secondary wall formation in fibers. Plant J. 63, 469–483 (2010).

42. S. Lohse, W. Schliemann, C. Ammer, J. Kopka, D. Strack., T. Fester, Organization and metabolism of plastids and mitochondria in arbuscular mycorrhizal roots of Medicago truncatula. Plant Physiol. 139, 329–340 (2005).

43. A. Liu, C.A. Contador, K. Fan., H.M. Lam, Interaction and regulation of carbon, nitrogen, and phosphorus metabolisms in root nodules of legumes. Front. Plant Sci. 9, 1860 (2018).

44. J.I. Sprent, Evolving ideas of legume evolution and diversity: a taxonomic perspective on the occurrence of nodulation. New Phytol. 174, 11–25 (2007).

45. M. Kamioka, S. Takao, T. Suzuki, K. Taki, T. Higashiyama, T. Kinoshita., N. Nakamichi, Direct repression of evening genes by CIRCADIAN CLOCK-ASSOCIATED1 in the Arabidopsis circadian clock. Plant Cell 28, 696–711 (2016).

46. C.E. Grant, T.L. Bailey., W.S. Noble, FIMO: scanning for occurrences of a given motif. Bioinformatics 27, 1017–1018 (2011).

47. J. Montiel, J.A. Downie, A. Farkas, P. Bihari, R. Herczeg, B. Balint, P. Mergaert, A. Kereszt., E. Kondorosi, Morphotype of bacteroids in different legumes correlates with the number and type of symbiotic NCR peptides. Proc. Natl. Acad. Sci. U.S.A. 114, 5041– 5046 (2017).

48. T.C. de Bang, P.K. Lundquist, X. Dai, C. Boschiero, Z. Zhuang, P. Pant, I. Torres-Jerez, S. Roy, J. Nogales, V. Veerappan, R. Dickstein, M.K. Udvardi, P.X. Zhao., W.R. Scheible, Genome-wide identification of Medicago peptides involved in macronutrient responses and nodulation. Plant Physiol. 175, 1669–1689 (2017).

49. P. Mergaert, A. Kereszt., E. Kondorosi, Gene expression in nitrogen-fixing symbiotic nodule cells in Medicago truncatula and other nodulating plants. Plant Cell 32, 42–68 (2020).

50. V. Dalla Via, C. Narduzzi, O.M. Aguilar, M.E. Zanetti., F.A. Blanco, Changes in the common bean transcriptome in response to secreted and surface signal molecules of Rhizobium etli. Plant Physiol. 169, 1356–1370 (2015).

51. Y. Kong, L. Han, X. Liu, H. Wang, L. Wen, X. Yu, X. Xu, F. Kong, C. Fu, K.S. Mysore, J. Wen., C. Zhou, The nodulation and nyctinastic leaf movement is orchestrated by clock gene LHY in Medicago truncatula. J Int. Plant Biol. 62, 1880–1895 (2020).

52. T. Mizoguchi, K. Wheatley, Y. Hanzawa, L. Wright, M. Mizoguchi, H.R. Song, I.A. Carre., G. Coupland, LHY and CCA1 are partially redundant genes required to maintain circadian rhythms in Arabidopsis. Dev. Cell 2, 629–641 (2002).

53. K.D. Edwards, J.R. Lynn, P. Gyula, F. Nagy., A.J. Millar, Natural allelic variation in the temperature-compensation mechanisms of the Arabidopsis thaliana circadian clock. Genetics 170, 387–400 (2005).

54. J. Ni, L. Dong, Z. Jiang, X. Yang, Z. Chen, Y. Wu., M. Xu, Comprehensive transcriptome analysis and flavonoid profiling of Ginkgo leaves reveals flavonoid content alterations in day-night cycles. PLoS One 13, e0193897 (2018).

55. F.D. Dakora, D.A. Phillips, Diverse functions of isoflavonoids in legumes transcend anti-microbial definitions of phytoalexins. Phys. and Mol. Plant Path. 49, 1–20 (1996).

56. N.K. Peters, J.W. Frost., S.R. Long, A plant flavone, luteolin, induces expression of Rhizobium meliloti nodulation genes. Science 233, 977–980 (1986).

57. J. Shin, K. Heidrich, A. Sanchez-Villarreal, J.E. Parker., S.J. Davis, TIME FOR COFFEE represses accumulation of the MYC2 transcription factor to provide time-of-day regulation of jasmonate signaling in Arabidopsis. Plant Cell 24, 2470–2482 (2012).

58. D.S. Chen, C.W. Liu, S. Roy, D. Cousins, N. Stacey., J.D. Murray, Identification of a core set of rhizobial infection genes using data from single cell-types. Front. Plant Sci. 6, 575 (2015).

59. C.W. Liu, J.D. Murray, The role of flavonoids in nodulation host-range specificity: an update. Plants (Basel) 5, (2016).

60. Q. Wang, S. Yang, J. Liu, K. Terecskei, E. Abraham, A. Gombar, A. Domonkos, A. Szucs, P. Kormoczi, T. Wang, L. Fodor, L. Mao, Z. Fei, E. Kondorosi, P. Kalo, A. Kereszt., H. Zhu, Host-secreted antimicrobial peptide enforces symbiotic selectivity in Medicago truncatula. Proc. Natl. Acad. Sci. U.S.A. 114, 6854–6859 (2017).

61. S. Yang, Q. Wang, E. Fedorova, J. Liu, Q. Qin, Q. Zheng, P.A. Price, H. Pan, D. Wang, J.S. Griffitts, T. Bisseling., H. Zhu, Microsymbiont discrimination mediated by a host-secreted peptide in Medicago truncatula. Proc. Natl. Acad. Sci. U.S.A. 114, 6848–6853 (2017).

62. R.C. O’Malley, S.C. Huang, L. Song, M.G. Lewsey, A. Bartlett, J.R. Nery, M. Galli, A. Gallavotti., J.R. Ecker, Cistrome and epicistrome features shape the regulatory DNA landscape. Cell 165, 1280–1292 (2016).

63. N. Nakamichi, Adaptation to the local environment by modifications of the photoperiod response in crops. Plant Cell Physiol. 56, 594–604 (2015).

64. M.W. Li, H.M. Lam, The modification of circadian clock components in soybean during domestication and improvement. Front Genet. 11, 571188 (2020).

65. M. Tadege, Wen J, He J, Tu H, Kwak Y, Eschstruth A, Cayrel A, Endre G, Zhao PX, Chabaud M, Ratet P, Mysore KS., Large scale insertional mutagenesis using Tnt1 retrotransposon in the model legume Medicago truncatula. Plant J. 54, 335–347 (2008).

66. T. Zielinski, A.M. Moore, E. Troup, K.J. Halliday., A.J. Millar, Strengths and limitations of period estimation methods for circadian data. PLoS One 9, e96462 (2014).

67. X. Cheng, M. Wang, H.K. Lee, M. Tadege, P. Ratet, M. Udvardi, K.S. Mysore., J. Wen, An efficient reverse genetics platform in the model legume Medicago truncatula. New Phytol. 201, 1065–1076 (2014).

68. V. Veerappan, K. Kadel, N. Alexis, A. Scott, I. Kryvoruchko, S. Sinharoy, M. Taylor, M. Udvardi., R. Dickstein, Keel petal incision: a simple and efficient method for genetic crossing in Medicago truncatula. Plant Methods 10, 11 (2014).

69. K.J. Livak, T.D. Schmittgen, Analysis of relative gene expression data using real-time quantitative PCR and the 2(-Delta Delta C(T)) Method. Methods 25, 402–408 (2001).

70. T.L. Bailey, J. Johnson, C.E. Grant., W.S. Noble, The MEME Suite. Nucleic Acids Res. 43, W39–49 (2015).

71. K. Katoh, D.M. Standley, MAFFT multiple sequence alignment software version 7:improvements in performance and usability. Mol. Biol. Evol. 30, 772–780 (2013).

72. A. Branca, T.D. Paape, P. Zhou, R. Briskine, A.D. Farmer, J. Mudge, A.K. Bharti, J.E. Woodward, G.D. May, L. Gentzbittel, C. Ben, R. Denny, M.J. Sadowsky, J. Ronfort, T. Bataillon, N.D. Young., P. Tiffin, Whole-genome nucleotide diversity, recombination, and linkage disequilibrium in the model legume Medicago truncatula. Proc. Natl. Acad. Sci. U.S.A. 108, E864–870 (2011).

73. D.W. Mount, Using gaps and gap penalties to optimize pairwise sequence alignments. CSH Protoc 2008, pdb top40 (2008).

74. S. Gupta, J.A. Stamatoyannopoulos, T.L. Bailey., W.S. Noble, Quantifying similarity between motifs. Genome Biol. 8, R24 (2007).

75. J.M. Franco-Zorrilla, I. Lopez-Vidriero, J.L. Carrasco, M. Godoy, P. Vera., R. Solano, DNA-binding specificities of plant transcription factors and their potential to define target genes. Proc. Natl. Acad. Sci. U.S.A. 111, 2367–2372 (2014).

76. S.F. Altschul, W. Gish, W. Miller, E.W. Myers., D.J. Lipman, Basic local alignment search tool. J. Mol. Biol. 215, 403–410 (1990).

77. S. Andrews, FastQC: a quality control tool for high throughput sequence data. Available online at: http://www.bioinformatics.babraham.ac.uk/projects/fastqc (2010).

78. A.M. Bolger, M. Lohse., B. Usadel, Trimmomatic: a flexible trimmer for Illumina sequence data. Bioinformatics 30, 2114–2120 (2014).

79. R. Patro, G. Duggal, M.I. Love, R.A. Irizarry., C. Kingsford, Salmon provides fast and bias-aware quantification of transcript expression. Nat Methods 14, 417–419 (2017).

80. R. Yang, Z. Su, Analyzing circadian expression data by harmonic regression based on autoregressive spectral estimation. Bioinformatics 26, i168–174 (2010).

81. D.T. Jones, W.R. Taylor., J.M. Thornton, The rapid generation of mutation data matrices from protein sequences. Comput. Appl. Biosci. 8, 275–282 (1992).

82. I. Letunic., P. Bork, Interactive Tree Of Life (iTOL) v4: recent updates and new developments. Nucleic Acids Res. 47, W256–W259 (2019).

